# A phenomenological model of neuroimmune interactions in epileptogenesis

**DOI:** 10.1101/2021.07.30.454477

**Authors:** Danylo Batulin, Fereshteh Lagzi, Annamaria Vezzani, Peter Jedlicka, Jochen Triesch

## Abstract

Epilepsy can have many different causes and its development (epileptogenesis) involves a bewildering complexity of interacting processes. Here, we present a first-of-its-kind computational model to better understand the role of neuroimmune interactions in the development of acquired epilepsy. Our model describes the interactions between neuroinflammation, blood-brain barrier disruption, neuronal loss, circuit remodeling, and seizures. Formulated as a system of nonlinear differential equations, the model is validated using data from animal models that mimic human epileptogenesis caused by infection, status epilepticus, and blood-brain barrier disruption. The mathematical model successfully explains characteristic features of epileptogenesis such as its paradoxically long timescales (up to decades) despite short and transient injuries, or its dependence on the intensity of an injury. Furthermore, stochasticity in the model captures the variability of epileptogenesis outcomes in individuals exposed to identical injury. Notably, in line with the concept of degeneracy, our simulations reveal multiple routes towards epileptogenesis with neuronal loss as a sufficient but non-necessary component. We show that our framework allows for *in silico* predictions of therapeutic strategies, providing information on injury-specific therapeutic targets and optimal time windows for intervention.

## Introduction

Epilepsy is a common neurological disorder that affects numerous physiological mechanisms in the central nervous system (***Lytton, 2008***; ***Devinsky et al., 2018***). Several processes play an important role in the development of epilepsy. These include the activation of innate and adaptive immune responses (***Ravizza et al., 2008***; ***Bauer et al., 2017***), disruption of the blood-brain barrier (BBB) integrity (***Marchi et al., 2012***; ***Löscher and Friedman, 2020***), neuronal loss (***Tasch et al., 1999***; ***Dingledine et al., 2014***), and remodeling of neural circuits (***Liao et al., 2011***; ***Bertram, 2013***; ***Jo et al., 2019***). Among the common causes of acquired epilepsy are traumatic brain injury (***Annegers and Coan, 2000***; ***Pitkänen and Immonen, 2014***), stroke (***Olsen, 2001***; ***Pitkänen et al., 2016***), central neural system infection (***Van Baalen et al., 2010***; ***Ramantani and Holthausen, 2017***), and new onset status epilepticus (SE) (***Holtkamp et al., 2005***; ***Gaspard et al., 2018***). All three lead to neuroinflammation, which has been widely studied in the context of epilepsy (***Barker-Haliski et al., 2017***; ***Rana and Musto, 2018***; ***Vezzani et al., 2019***). In particular, signaling pathways of neuroinflammation have been shown to modulate neuronal excitability (***Devinsky et al., 2013***; ***Vezzani and Viviani, 2015***), seizure threshold (***Heida et al., 2004***; ***Galic et al., 2008***) and severity of the seizure burden (***Auvin et al., 2010b***; ***Tan et al., 2015***; ***Rana and Musto, 2018***). For example, ***Patel et al***. (***2017***) have shown that TNF knockout mice have a significantly reduced seizure burden. A similar effect was observed in animals with a deletion of TNFR1, while ablation of TNFR2 led to an increase of seizure burden. Moreover, it has been hypothesized that noxious stimuli causing neuroinflammation may not only be associated with an acute injury but also with abnormal neuronal activity, resulting in so-called neurogenic neuroinflammation (***Xanthos and Sandkühler, 2014***).

The complexity of the involved processes interacting on various timescales makes understanding epileptogenesis (EPG) a great challenge. In such a complex system with several nonlinear interacting processes, mathematical modeling is a useful tool for a better understanding the system’s dynamics. In addition, modeling helps to systematize and explain the great body of observations obtained in clinical and animal model studies, and to provide valuable predictions for further experimental studies. The mathematical modeling approach has already been successfully applied to studying the dynamics of ictogenesis (initiation, spreading, and termination of epileptic seizures) (***Jirsa et al., 2014***; ***Proix et al., 2017***; ***Jirsa et al., 2017***). However, no computational model of EPG that implements multiple major pathomechanisms at their relevant time scales (including neuroinflammation) has yet been developed.

Therefore, in this paper, we present a first-of-its-kind mathematical model that describes and simulates the course of acquired epilepsy development and accounts for the epileptogenic role of neuroimmune interactions. Being tested on data from three animal models, our unified computational framework allows for simulation of EPG caused by most common types of neural injuries using a single parameter set. It explains, among other things, how different causes (injury types) lead to similar outcomes (epilepsies) and why the development of epilepsy may take months or years after the triggering injury in certain conditions. Our mathematical analysis reveals that these long time scales result from the underlying dynamical system slowing down and lingering in the vicinity of an unstable fixed point. In addition to the reproduction and explanation of injury-specific features of EPG, our model provides insights into general principles of EPG and allows us to generate testable predictions for different therapeutic strategies.

## Methods

### Model structure and interactions captured by the model

The model describes interactions between neuroinflammation (*I*), BBB disruption (*B*), neuronal loss (*D*), circuit remodeling (*R*) and epileptic seizures (*S*) upon neurological injury (Fig. 1A). The probability of spontaneous recurrent seizure occurrence is assumed to depend on two seizure-promoting factors: intensity of neuroinflammation and degree of pathological circuit remodeling (arrows *I* → *S*, *R* → *S* in Fig. 1A). Pathological remodeling in circuits (*R*) may be constituted by, among others, loss of inhibitory neurons (***Sloviter, 1987***; ***Knopp et al., 2008***), abnormal excitatory synaptogenesis (***Weissberg et al., 2015***; ***Kim et al., 2017b***), or increase of recurrency in neural circuits due to mossy fiber sprouting (***Tauck and Nadler, 1985***; ***Buckmaster, 2014***). Such pathological remodeling leads to an increase of excitability in neuronal circuits and rising of the chance of critical synchronization that results in the occurrence of spontaneous seizures (arrow *R* → *S*). A facilitation of a neuroinflammatory response was shown to modulate neuronal excitability (***Devinsky et al., 2013***; ***Vezzani and Viviani, 2015***) and, consequently, lower seizure threshold (***Heida et al., 2004***; ***Galic et al., 2008***) (arrow *I* → *S*). In agreement with a recent finding of the inhibition of neuronal activity by microglia (***Badimon et al., 2020***), we assume that in conditions of profound neuroinflammation this physiological function may be impaired due to adoption of proinflammatory phenotypes by microglia.

**Figure 1.**
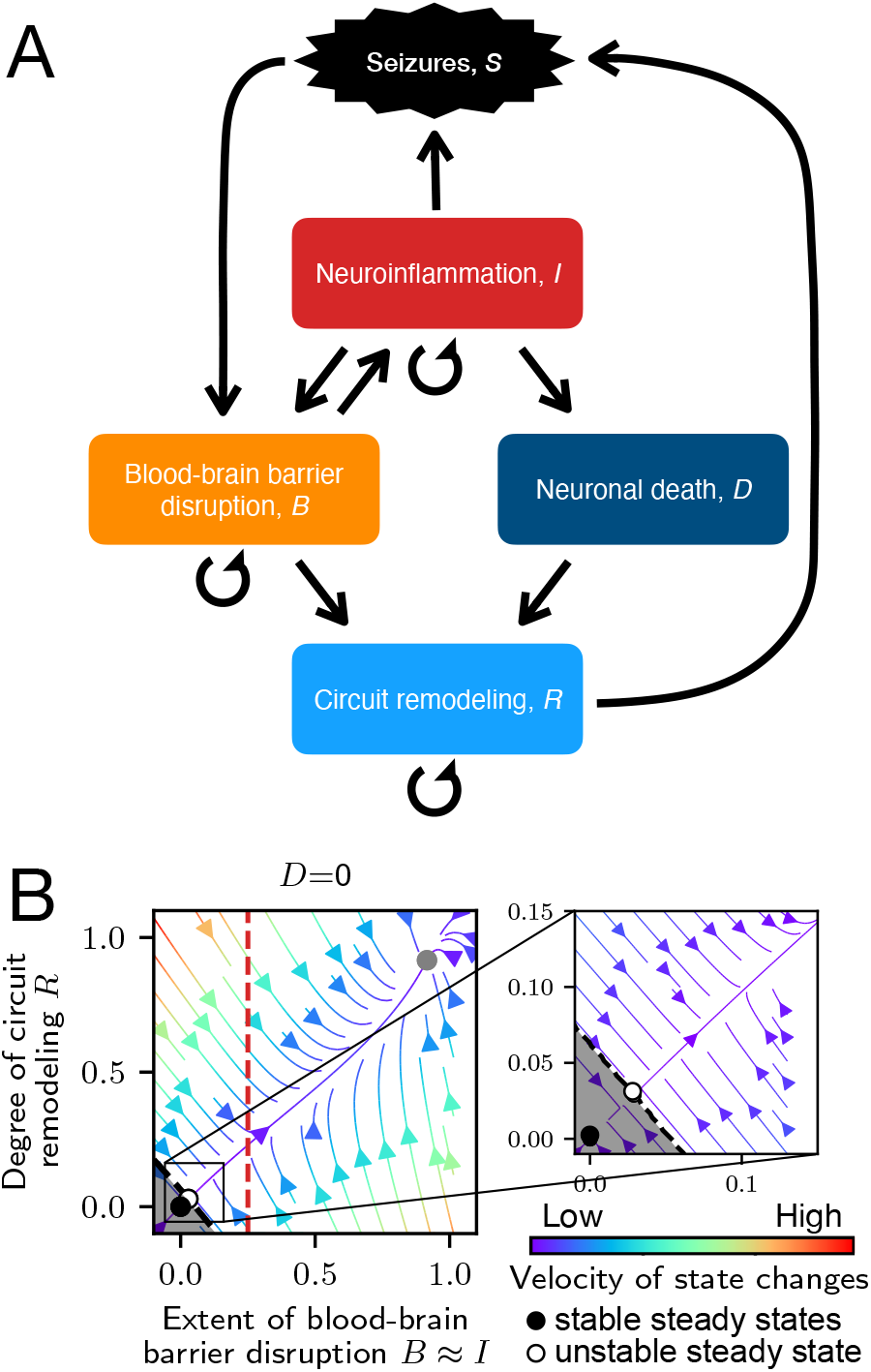
Model overview: **A**. Interactions between variables in our model. **B**. State space plot for the rate model with 3 steady states: ‘healthy’ stable steady state (black), unstable fixed point (white), and ‘epileptic’ stable steady state corresponding to state of progressed EPG (gray). The dashed black line going through the unstable fixed point is a separatrix, which separates the basin of attraction of the ‘healthy’ steady state (shaded area) from the basin of attraction of the ‘epileptic’ steady state. Color of arrows indicates the velocity of state changes. Red dashed line corresponds to the glial neurotoxicity threshold Θ under conditions of timescales separation *I* ≈ *B*.

Epileptic seizures have been shown to cause metabolic stress on the central nervous system due to increased energy demands associated with excessive neural activity (***Zhang et al., 2015***; ***Prager et al., 2019***). Data from humans and animal models also suggest that seizures induce leakiness of the BBB and that the leakiness is negatively correlated with time since the last seizure (***Van Vliet et al., 2007***; ***Rüber et al., 2018***). In our model, we account for this effect of seizures (Fig. 1A, arrow *S* → *B*), as well as neuroinflammation that can be caused by exposure of the parenchyma to cells or soluble factors infiltrating through the leaky BBB (***Farrell et al., 2017***; ***Löscher and Friedman, 2020***) (arrow *B* → *I*). Neuroinflammation itself can cause leakiness of the BBB via activation of cells of the neurovascular unit (***Obermeier et al., 2013***), which results in a positive feedback loop in our model (Fig. 1A, arrow *I* → *B*). Excessive neuroinflammation may lead to neurotoxicity (***Block et al., 2007***; ***Biber et al., 2014***), which is also accounted for in our model (arrow *I* → *D*). Neural loss, in turn, leads to remodeling of neural circuits. Some remodeling may aim at maintaining functional properties. Here we only consider the kind of remodeling that leads to increased seizure susceptibility (arrow *D* → *R*). Furthermore, we also account for pathological circuit remodeling that is independent of neural loss. Examples of such remodeling are development of chronic inhibition deficits (***Kim et al., 2017b***) and excessive excitatory synaptogenesis (***Weissberg et al., 2015***) due to albumin extravasation upon BBB dysfunction (Fig. 1A, arrow *B* → *R*).

### Mathematical description

The interactions between the processes described above are modeled with a system of stochastic nonlinear ordinary differential equations:

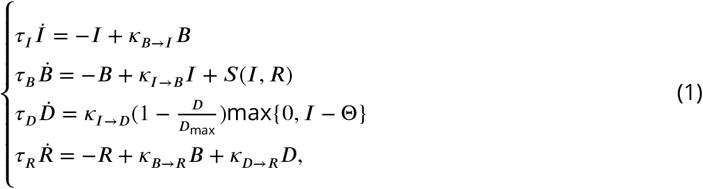

where *I*(*t*) is neuroinflammation intensity; *B*(*t*) is the extent of BBB disruption; *D*(*t*) is the extent of neuronal loss; *R*(*t*) is the degree of circuit remodeling. All variables are assumed to be reversible, except for neural loss *D*(*t*), which is motivated by the impossibility of recovery of dead neurons and thereby excludes the of possibility of neurogenesis in the adult nervous system. *κ*_*B*→*I*_, *κ*_*I*→*B*_, *κ*_*I*→*D*_, *κ*_*B*→*R*_, *κ*_*D*→*R*_ are parameters for coupling strengths of the respective variables. The described processes are assumed to operate on 3 timescales: fast (seconds-minutes) for epileptic seizures; intermediate (hours-days) for neuroinflammatory reaction (*τ*_*I*_); and slow (days-weeks) for permeability of the BBB (*τ*_*B*_), neuronal loss (*τ*_*D*_) and circuit remodeling (*τ*_*R*_).

Mild neuroinflammation, which is a physiological process aiming to maintain tissue homeostasis (***Yong et al., 2019***), is assumed to have no neurotoxic effects. Thus, neurotoxicity leading to neuronal loss requires excessively activated glia (*I*(*t*) > Θ), where Θ is a neurotoxicity threshold and *D*_max_ is a maximum possible extent of neuronal loss.

The occurrence of spontaneous recurrent seizures in the system is modeled with a Poisson process. The seizure rate (the probability of a seizure occurring per unit time) is monotonically increasing with the intensity of neuroinflammation *I*(*t*) and the extent of circuit remodeling *R*(*t*) according to:

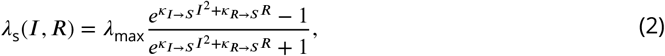

where *λ*_max_ is a maximum possible amount of seizures per day; *κ*_*I*→*S*_ and *κ*_*R*→*S*_ are parameters scaling the seizure-promoting contribution of, respectively, neuroinflammation and circuit remodeling. The sigmoid shape of the function (Appendix 1) reflects the saturation effect of maximum possible seizure burden that nervous system may be exposed to within a finite time interval due to metabolic constraints. The assumption of a quadratic dependence of the seizure rate on the intensity of neuroinflammation minimizes seizure-promoting effects of mild neuroinflammation *I*(*t*) ≳ 0.

The term *S*(*I, R*) in Eq. 1 describes an effect of seizure activity on permeability of the BBB:

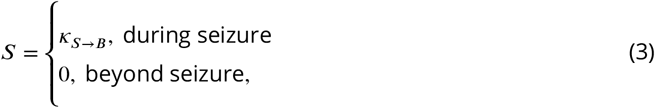

where 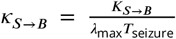 describes the effect of a single seizure on the permeability of the BBB; *K*_*S*→*B*_ is a parameter defining the maximum possible burden of seizure activity; *T*_seizure_ is the seizure duration.

### Time scale separation and rate model

In addition to the model simulating stochastically occurring seizures (Eqs. 1–3), we developed a rate model, where the Poisson process is approximated with a seizure burden function:

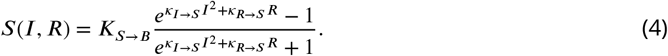

The rate of seizures dictated by Eq. 4 does not allow for tracking the occurrence of individual seizures, but it provides a means for more intuitive explanation of the dynamics of the system. Moreover, we can perform the time scale separation procedure for the equation describing neuroinflammation since its timescale is smaller than the timescales of the other processes (*τ_I_* < *τ_B_*, *τ_D_*, *τ_R_*). After performing the time scale separation (see Appendix 2 for details), the fast evolution of the neuroinflammation variable can be approximated by the dynamics of BBB disruption variable according to Eq. 1: *I*(*t*) ≈ *B*(*t*). Thus, we can obtain the state space representations of the model in the *B* − *R* domain for particular values of the monotonically rising variable *D*(*t*). Stability analysis (Appendix 3) shows that in the absence of neuronal loss, the system is bistable, having 3 steady states: a ‘healthy’ steady state in the origin; an unstable fixed point; and a stable steady state corresponding to the state of progressed EPG (Fig. 1B). A separatrix, illustrated with a black dashed line, passes through the unstable fixed point and separates the basins of attraction of the two stable steady states. The neurotoxicity threshold (for neuronal death), illustrated with the red dashed line, divides the state space into two areas: to the left are states in which no neuronal loss is being induced, and to the right are states in which the neuronal population experiences neurotoxic effects due to glial overactivation.

### Simulation of various neurological injuries

Using a single set of parameters (Appendix 4), the mathematical model allows for the simulation of EPG progression caused by various types of neurological injuries. Modeling of different types of injuries is carried out by application of time sequences of perturbations mimicking pathological effects of the respective injury (Fig. 2A-C). In this study, we present simulation of three injury types and associated animal models, which are distinct in methods of EPG induction, and time course of disease pathology. Detailed simulation protocols are available in Appendix 5.

**Figure 2.**
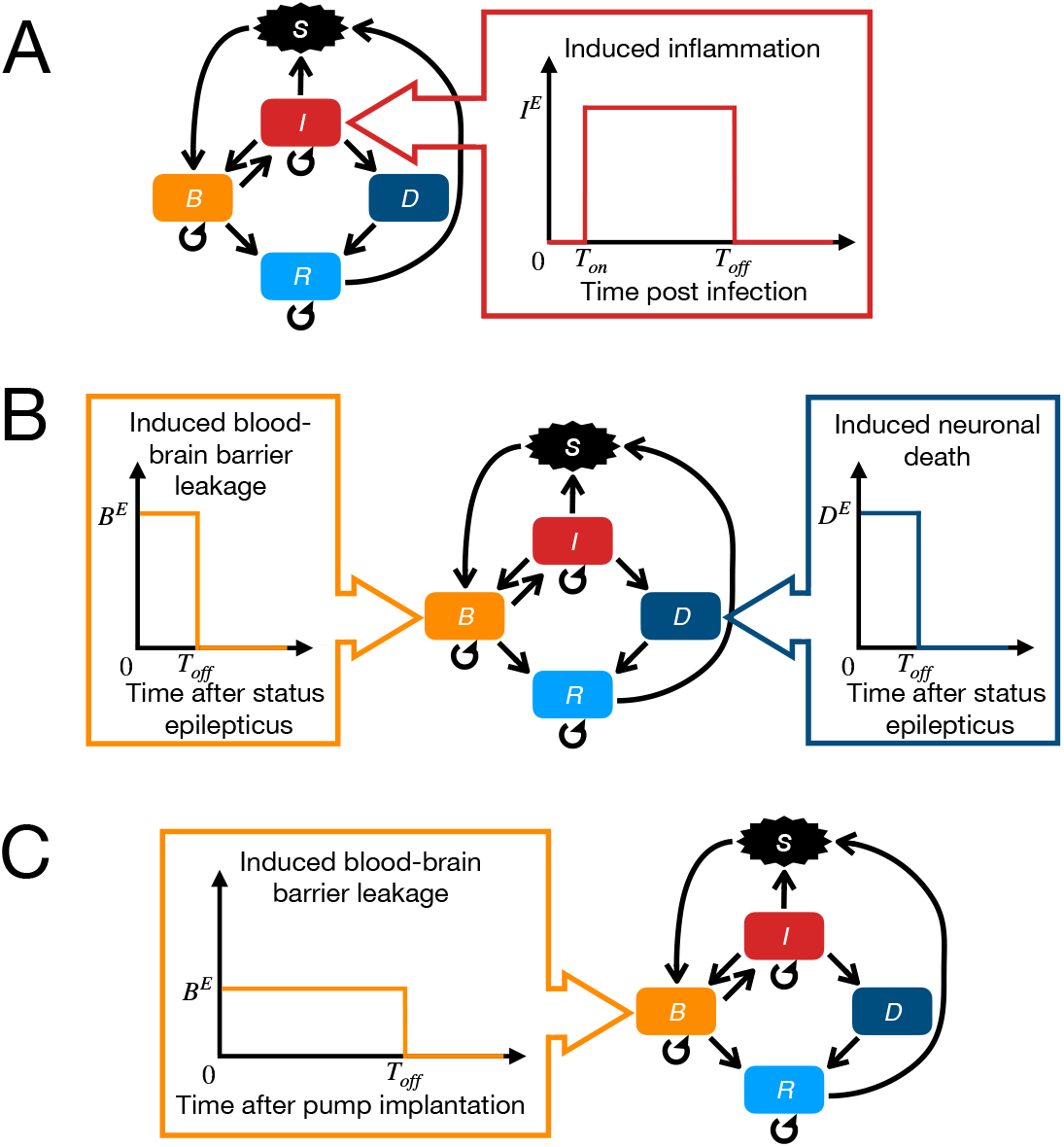
Simulation schematics for three animal models of epileptogenesis: **A**. Theiler’s murine encephalomyelitis virus (TMEV) mouse model. **B**. Chemically-induced (pilocarpine) SE rodent model. **C**. BBB disruption rodent model.

The first animal model is Theiler’s murine encephalomyelitis virus (TMEV) mouse model, which is commonly used in epilepsy research for modeling infection-induced epilepsy in humans (***Libbey et al., 2008***; ***Stewart et al., 2010***; ***Libbey et al., 2011***; ***Gerhauser et al., 2019***). During the first week post infection, the acute neuroinflammatory response is developing. It is characterized by the presence of excessive concentrations of proinflammatory molecules (***Patel et al., 2017***), micro- and astrogliosis (***Kirkman et al., 2010***), which are followed by relative recovery during subsequent weeks post infection. Taking this dynamics of pathology development into account, we simulate infectious injury by induction of neuroinflammation with onset and offset within the first week post infection (Fig. 2A).

The second animal model is a pilocarpine rodent model, which is based on induction of SE (excessively long generalized seizure) by injection of pilocarpine with further pharmacological termination of SE (***Polascheck et al., 2010***; ***Zhang et al., 2015***; ***Brackhan et al., 2016***; ***Kim et al., 2017a***). SE induces neuronal loss, neuroinflammation and profound leakage of the BBB. In our framework, we do not define the inflammatory perturbation for simulation of pilocarpine since it would be indirectly induced by BBB disruption. Thus, we simulate the induction of pilocarpine model injury as a combination of two external perturbations (Fig. 2B): neuronal death, which allows us to account for initial SE-associated neuronal loss (***Auvin et al., 2010a***), and BBB leakage, which normalizes after 1-2 days post SE (***Bankstahl et al., 2018***).

The third animal model is based on the induction of BBB leakage. It is obtained via exposure of the brain tissue to an artificial cerebrospinal fluid containing bile salts (***Seiffert et al., 2004***; ***Tomkins et al., 2007***). Alternatively, an artificial cerebrospinal fluid may contain serum albumin, which mimics the extravasation of this protein in brain parenchyma when blood-brain barrier is dysfunctional (***Weissberg et al., 2015***). This animal model is simulated by setting *B* to a high value for a period corresponding to the time of application of bile salts or pumping albumin into the brain of an animal (Fig. 2C).

### Statistics

All experiments were performed as mathematical model simulations. Group allocation of samples is described in Appendix 5. No data were excluded as outliers. For stochastic model simulations, the sample size of N=30 was chosen, which is twice larger than a common sample size in animal model experiments (***Weissberg et al., 2015***; ***Patel et al., 2017***; ***Kirkman et al., 2010***; ***Brackhan et al., 2016***; ***Zhang et al., 2015***). Data are presented both in raw format (e.g. neuroinflammation intensity development over time, time sequences of seizures occurrence), and in format of mean ± standard error of the mean (SEM), when convenient (e.g. latent period duration, seizure occurrence frequency for the whole sample). Due to bistability of system states (Fig. 1B, Appendix 3), data could not be assumed to have Gaussian statistics and non-parametric tests were used. Specifically, an unpaired two-group Mann-Whitney U test was performed for analyses of dose-dependence effects on the intensity of the injury. *p* = 0.05 was chosen as a threshold for statistical significance and exact p-values were reported.

### Data and code availability

Experimental data from animal models of epileptogenesis, which are used in this study, are described in detail in Appendix 6. Simulation code, data, analysis and figures production scripts are available at https://github.com/danylodanylo/math-model-epileptogenesis.git.

## Results

### Dependence of EPG on injury intensity and emergence of long timescales

The risk of epilepsy development and severity of seizure burden have been shown to depend on the intensity of the neurological injury. Patients suffering from mild traumatic brain injury (TBI) have 2.1% cumulative probability of seizure development over 30 years, while for severe TBI it rises up to 16.7% (***Annegers et al., 1998***). The dose-dependence is not specific to TBI and also translates to animal models of epilepsy. In infection-induced epilepsy in mice, increasing viral dose from 3 × 10^3^ plaque forming units (PFU) to 3×10^6^ PFU leads to an increase of the fraction of animals with seizures from 25% to 80% (***Libbey et al., 2011***). In SE models, the duration of the acute epileptiform activity determines the incidence of epilepsy in animals (***Löscher, 2015***).

We tested whether our model captures the observed spectrum of dose-dependence effects of the risk of EPG (and its characteristic features) on the intensity of neurological injury. Figure 3 illustrates the results of simulation of EPG induced by BBB disruption. The latent period duration (time period between injury and arrival of first seizure, Fig. 3A) and seizure burden (Fig. 3B) in simulated animals are in agreement with data reported in the animal model study (***Weissberg et al., 2015***). Accordingly, seizure manifestation was observed in our simulations either during the infusion of albumin (7 days after pump implantation), or after albumin pump removal (e.g. animals #6, #7, #8 in Fig. 3C). Similarly to the animal model data, no neuronal death was observed within the observation period because neuroinflammation did not reach the neurotoxicity threshold for neuronal death (Fig. 3D).

**Figure 3.**
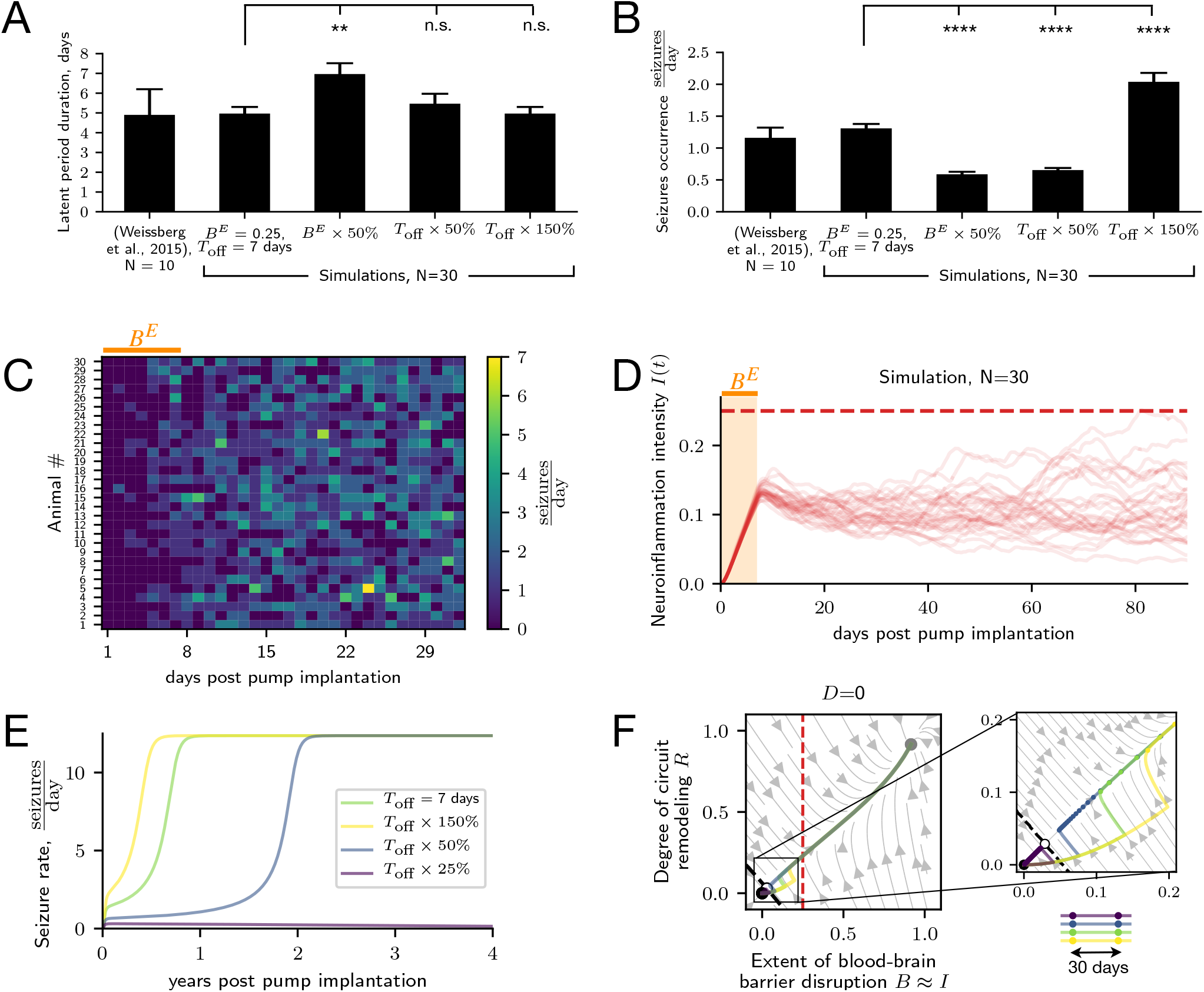
Dose-dependence of EPG on the intensity of neurological injury: **A**. Comparison of the latent period duration in BBB disruption animal model data (***Weissberg et al., 2015***) and simulations with the intensity of the injury: matched to ***Weissberg et al***. (***2015***) (*B^E^* = 0.25, *T*_off_ = 7 days); decreased via lowering albumin concentration (*B^E^* × 50%); decreased via shortening the time window of albumin infusion (*T*_off_ × 50%); increased via prolongation of the time window of albumin infusion (*T*_off_ × 150%). The *p* values for two-sided Mann-Whitney U test are respectively 0.0059, 0.7772, and 0.9939. **B**. Comparison of seizure burden on the first month after injury onset in BBB disruption animal model data (***Weissberg et al., 2015***) and simulations (annotation identical to caption in A.). The *p* values for two-sided Mann-Whitney U test are respectively 1.0662 10^−10^, 5.6488 10^−10^, and 1.1774 10^−8^. **C**. Time sequences of seizure occurrence in individual animals. Orange bar corresponds to the time window of injury induction. **D**. Time course of neuroinflammation in individual animals (N=30). Red dashed line corresponds to the neurotoxicity threshold Θ. Light orange area corresponds to the time window of injury induction. **E**. EPG in response to injuries of 4 different intensities illustrated with seizure rate development over time post pump implantation. Simulation results obtained with the rate model. The injury intensity control was implemented by modification of the duration of the time window of albumin infusion (*T*_off_). **F**. EPG in response to injuries of 4 different intensities (annotation identical to caption E.) illustrated over the state space plot. The state space consists of 3 steady states: ‘healthy’ (black), unstable (white) and ‘epileptic’ (gray). Dashed black line (separatrix) separates basins of attraction of two stable steady states. Red dashed line corresponds to neurotoxicity threshold Θ. Distance between circle markers on EPG traces correspond to time intervals of 30 days.

Our model predicts that with a 50% decrease in the injury intensity the latent period duration has prolonged (5.57 ± 0.34 days vs 7.23 ± 0.47 days, Fig. 3A), and the seizure burden has dropped significantly (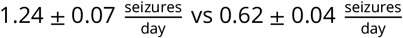, Fig. 3B). In the experimental setting, the manipulation of injury intensity used in this simulation corresponds to lowering of the albumin concentration in the infused artificial cerebrospinal fluid. An alternative approach to manipulation of injury intensity is prolongation or shortening of the time window of artificial cerebrospinal fluid infusion. Shortening by 50% did not have a significant effect on latent period duration (Fig. 3A), but led to a significant drop in the seizure burden (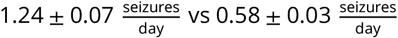, Fig. 3B). On the other hand, the increase of injury intensity simulated by 50% prolongation of infusion time window led to a significant rise of seizure burden (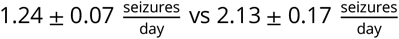, Fig. 3B).

In addition to latent period duration and seizure burden, the intensity of the injury has been also shown to affect the risk of epilepsy development itself. Figures 3E,F show that injury of low intensity does not lead to progressive EPG due to the restoration of the healthy state. Mathematically, the dose-dependence originates from the fact that injuries of low intensity fail to push the system state across the separatrix into the basin of attraction of the ‘epileptic’ steady-state (Fig. 3F). Instead, the model recovers without progressive EPG.

Surprisingly, but in line with clinical observations, remarkably long timescales of EPG (up to decades) emerge in our mathematical model, despite the ‘slowest’ variables operating on relatively fast timescales of weeks. From the mathematical point of view, slowing down of dynamics is occurring in our model when the state of the system approaches the unstable steady state (Fig. 1B). In order to visualize this property, we have performed a simulation of the injury with lowered intensity, which leads to slower EPG (Fig. 3E). In this case, the progression of the pathology takes longer due to slowing down of state changes around the unstable fixed point (Fig. 3F), while EPG will be facilitated when caused by injury of increased intensity. In sum, consistent with clinical data, our model can capture the effect of the intensity of the injury on the latent period duration, seizure burden, and the risk of EPG.

### Variability of EPG risk and pathology severity

In addition to the dose-dependent effects of injury intensity, the variability of EPG risk and the severity of pathology is evident even in animals exposed to identical injury. Even in highly standardized conditions of animal experiments, the fraction of animals that develop seizures, and the seizure burden in seizing animals are varying despite identical parameters of induced injury. For example, according to the data from a study by ***Polascheck et al***. (***2010***), in 12 rats treated with pilocarpine, 2 have not shown any seizures, while seizure frequency ranged from 1 to 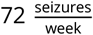 in the remaining animals at the time point of 8 weeks after the SE. The variability in EPG outcome is likely to depend on, among others, variability in genetic and epigenetic features of animals in experiment, and variability in various factors when conducting the experimental procedures. Moreover, the brain is an intrinsically stochastic complex system (***Deco et al., 2009***). Therefore, we added this intrinsic stochasticity to our model allowing for the variability in outcomes of EPG even in identical animals exposed to identical injuries.

To test whether our model is able to account for the experimentally observed variability of EPG outcomes, we simulated infection-induced EPG in the TMEV model (Fig. 4). Our simulation reproduces the characteristic temporal pattern where seizures manifest after second day post infection (day 2.83 ± 0.13) and profoundly drop in frequency during the second week post infection (Fig. 4A,F). Moreover, the computational model captures the characteristic time course of neuroinflammation (Fig. 4B) as well as neuronal death, which is characterized by the occurrence of macroscopically measurable neuronal damage on day 4 post infection and its further progression and saturation from the second week post infection (Fig. 4C). Our results suggest that this characteristic plateau of neuronal loss progression originates from the attenuation of the neuroinflammatory response during week 2 post infection (Fig. 4D). ***Kirkman et al***. (***2010***) have shown that neuronal death was significantly more abundant in animals that developed seizures versus those that did not. Consistent with this observation, neuronal loss in the model is correlated with seizure burden during the acute post infection stage (Fig. 4E).

**Figure 4.**
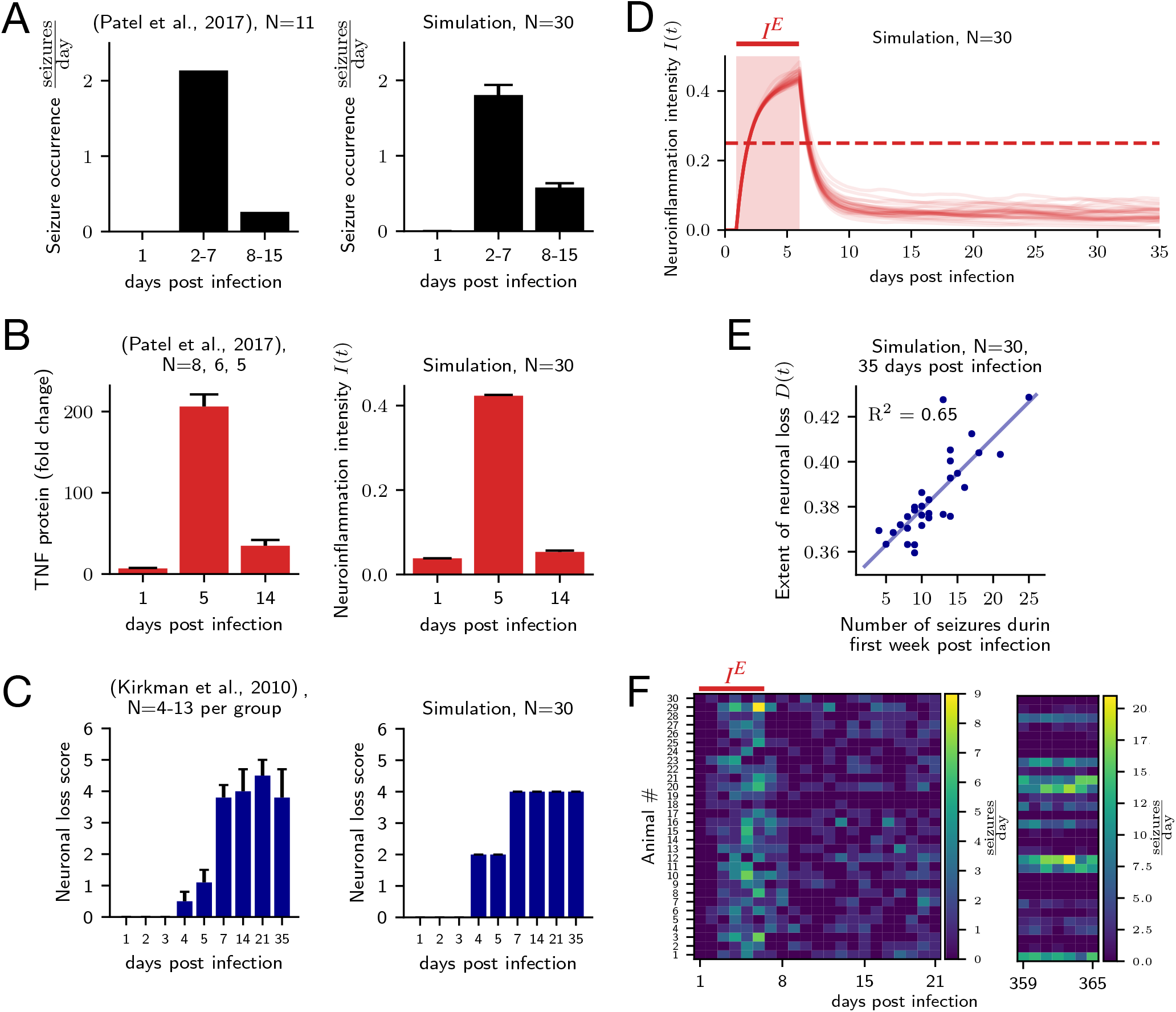
Model explains key features of infection-induced EPG: **A**. Comparison of characteristic seizure occurrence patterns from TMEV animal model data (left, ***Patel et al***. (***2017***)) and simulation (right). ***Patel et al***. (***2017***) reported the total number of seizures per day aggregated over N=11 animals together. **B**. Comparison of neuroinflammation time courses from TMEV model (left, ***Patel et al***. (***2017***)) and simulation (right). **C**. Comparison of neuronal loss score progression from TMEV model (left, ***Kirkman et al***. (***2010***)) and simulation (right). Neuronal loss score for the simulation was computed using the masking procedure from (***Kirkman et al., 2010***). Masking procedure and its effect of ‘masking out’ variability in the simulation results are explained in detail in the supplementary figure. **D**. Neuroinflammation course in individual animals (N=30). Red dashed line corresponds to the neurotoxicity threshold Θ. Light red area corresponds to the time window of injury induction. **E**. Neuronal loss one month post infection (day 35) is correlated with severity of seizure burden in the acute phase (week 1 post infection). Blue dots correspond to individual animals. Blue line corresponds to linear regression fit with coefficient of determination *R*^2^ = 0.65. **F**. Time sequences of seizure occurrence in individual animals. Red bar corresponds to the time window of injury induction. **Figure 4–Figure supplement 1. Neuronal loss score computation (masking procedure) from** *Kirkman et al*. (*2010*): Raw neuronal death data from TMEV model simulation (left) and neuronal loss score computation scheme (right). Horizontal dashed lines on the left correspond to 10%, 30% and 60% extent of neuronal loss, which are the border values separating score values in the scheme from *Kirkman et al*. (*2010*). In *Kirkman et al.* (*2010*), neuronal loss score data are presented as a sum of scores for 2 hippocampi (maximum score: 3 × 2 = 6). Thus, neuronal loss score computed for simulated TMEV animals was multiplied by factor of 2 for comparability with experimental data. Absence of variability (0 SEM) in Fig. 4C is explained by ‘masking out’ of variability in neuronal loss score computation (left).

Evaluation of seizure occurrence on a follow-up period of one year post infection (Fig. 4F, Fig. 5A,B) illustrates the variability of EPG outcomes in simulated animals. Among 30 simulated animals, 9 did not exhibit any seizures within one week, while the remaining 21 exhibited seizure burdens of various severity. Our model suggests that even for (hypothetically) identical animals exposed to identical injury, the EPG outcome is variable due to the stochastic nature of spontaneous recurrent seizures. Mathematically, the stochastic nature of seizure generation induces noise in the EPG (Fig. 5C), affecting disease progression and outcome (Fig. 5A,B).

**Figure 5.**
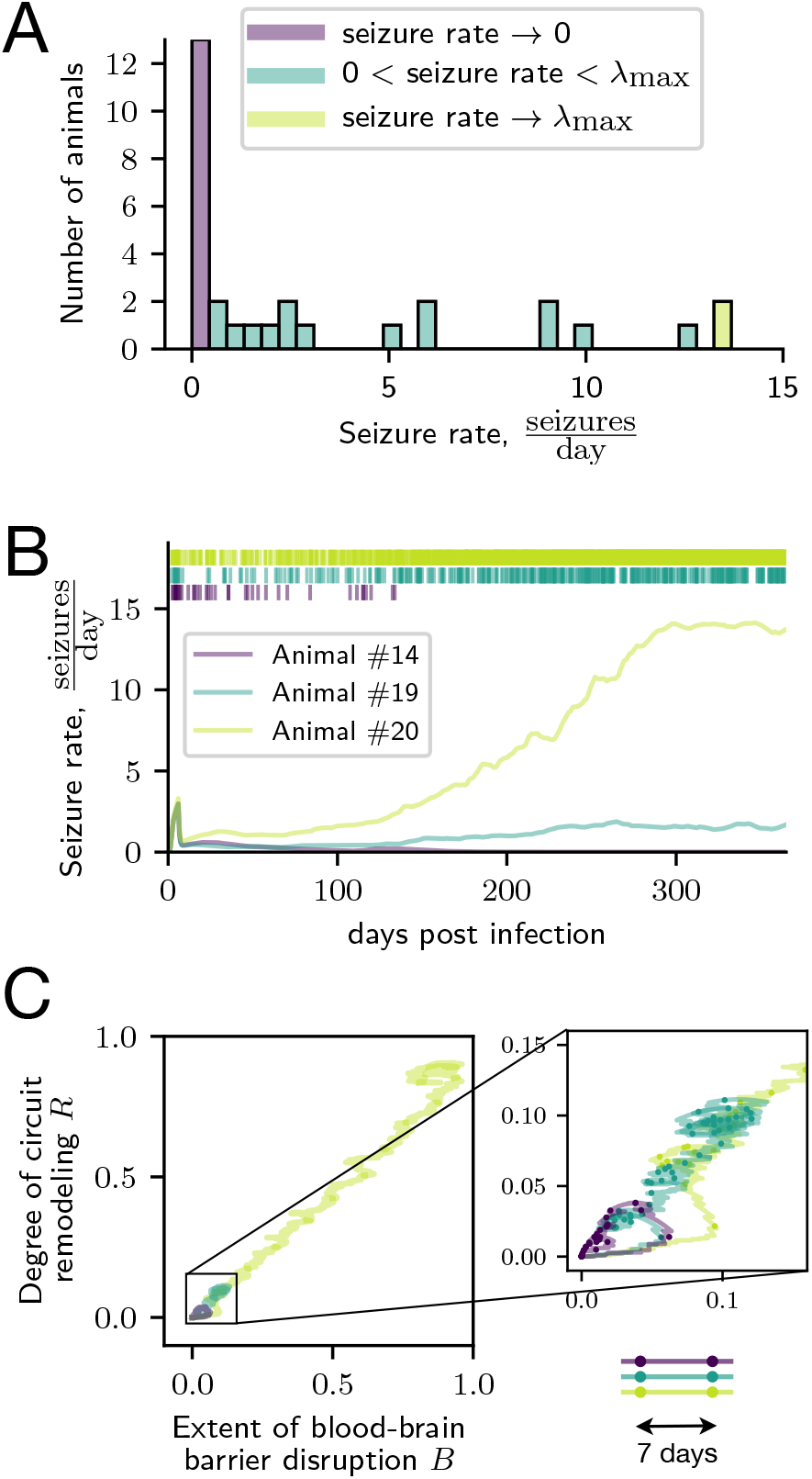
Variability of EPG outcomes in identical animals exposed to identical injury originates from the stochastic nature of spontaneous seizures: **A**. Distribution of seizure rate one year after infection for 30 simulated TMEV animals. **B**. Examples of seizure rate development in time for 3 animals with different seizure burden outcomes on one year post infection (line color code for 3 animals is consistent in all subfigures). The raster plot on top illustrates the occurrence of seizures in time for corresponding animals. **C**. EPG course for 3 animals with different seizure burden outcomes on one year post infection illustrated in B-R domain. Distance between circle markers on EPG traces correspond to time intervals of 7 days. Overall visualization period is 1 year post infection.

### Computational model accounts for complex and injury-specific features of EPG

EPG is often conceptualized as a two-stage process comprising a clinically silent latent period before the occurrence of a first spontaneous seizure and a subsequent seizure period. However, a growing body of evidence suggests that this view may be overly simplistic (***Pitkänen et al., 2015***). For example, in a perforant path stimulation rat epilepsy model epileptiform activity is already observed just after the injury induction prior to the first seizure (***Bumanglag and Sloviter, 2018***), questioning the existence of a latent period. Indeed, EPG appears to exhibit various injury-specific features. In the following, we show how our model can account for qualitatively different types of EPG. We illustrate this for different types of EPG-inducing injuries including BBB leakage, infection, and SE (due to pilocarpine administration) in Figs. 3, 4 and 6, respectively. In the case of EPG induced by BBB leakage, the latent period approaches one week in duration (4.97 ± 0.33 days, Fig. 3A,C). On the contrary, in the TMEV infection model, the occurrence of first spontaneous seizures takes place already after the second day after the viral infection (2.83 ± 0.13 days). Then, this early onset of seizures is followed by a period of profound decrease in seizure activity starting during the 2nd week post-infection (Fig. 4A,F).

**Figure 6.**
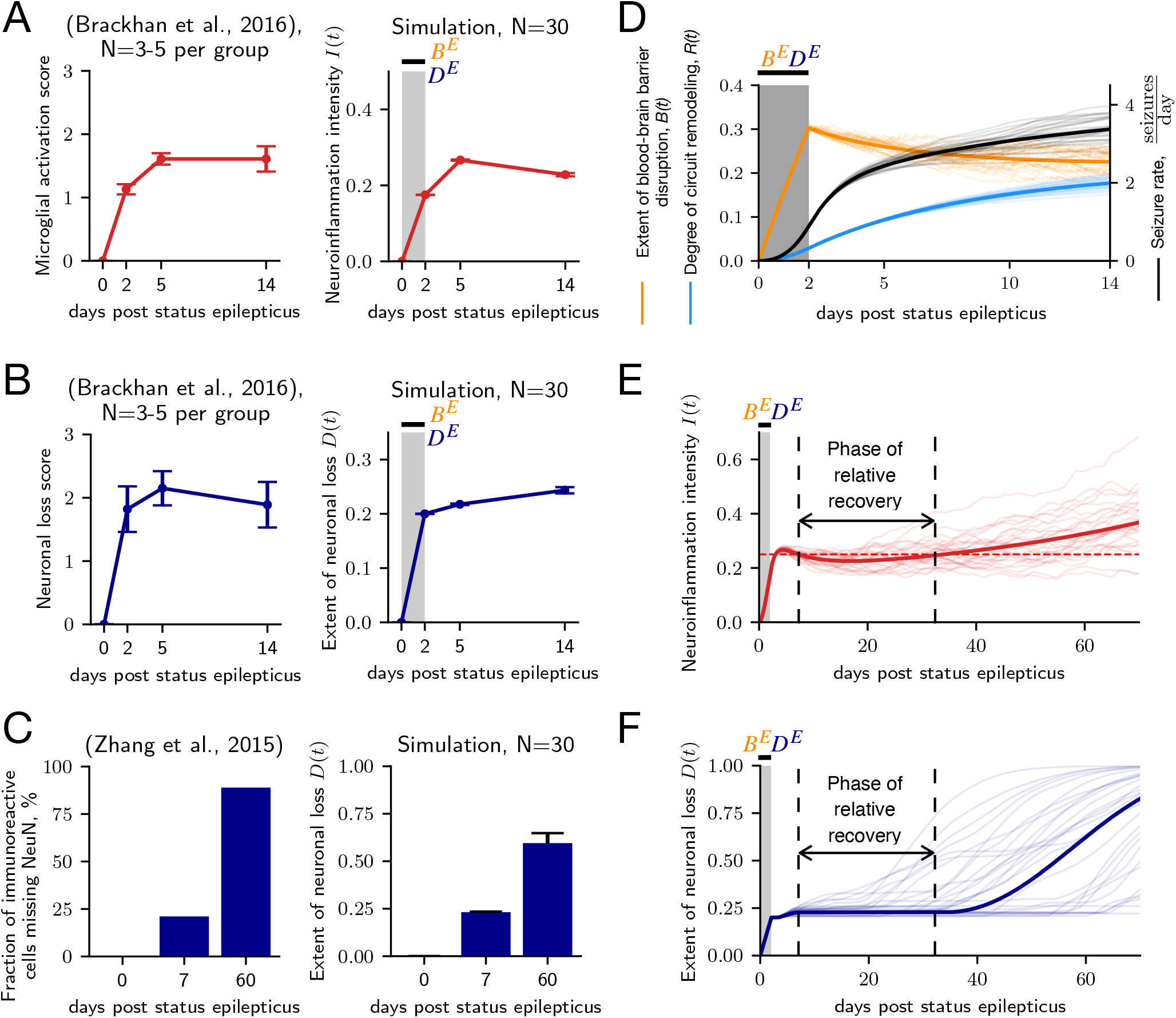
Mathematical model captures and explains mechanisms of injury-specific features of EPG in pilocarpine-induced SE animal model of epilepsy: **A**. Comparison of microglial activation progression from animal model (left, *Brackhan et al*. (*2016*)) with neuroinflammation course in simulated animals (right). Data are shown with mean values (red dots) and error bars for SEM. Gray area corresponds to the time window of injury induction. **B**. Comparison of neuronal loss progression from animal model (left, *Patel et al*. (*2017*)) with neuronal loss in simulated animals (right). Data are shown with mean values (blue dots) and SEM bars. Gray area corresponds to the time window of injury induction. **C**. Comparison of neuronal loss progression from animal model data (left, ***Zhang et al***. (***2015***)) with neuronal loss in simulated animals (right) illustrated for 3 time points. For simulation results, data are shown with mean values (blue bars) and error bars for SEM. **D**. Simulation results illustrate processes underlying the rise of seizure rate after injury despite relative recovery of the BBB integrity. Orange, light blue and black thin lines correspond respectively to extent of BBB disruption, degree of circuit remodeling and seizure rate in individual animals (N=30). Solid lines correspond to prediction from the rate model. Gray area corresponds to the time window of injury induction. **E**. Neuroinflammation development in time with indication of presumed phase of relative recovery characterized by absence of neurotoxicity in rate model prediction. Thin lines correspond to individual animals (N=30). Solid lines correspond to prediction from the rate model. Red dashed line corresponds to the neurotoxicity threshold Θ. Gray area corresponds to the time window of injury induction. **F**. Extent of neuronal loss development in time with indication of presumed phase of relative recovery characterized by absence of neurotoxicity in rate model prediction (annotation identical to caption in E.).

In the pilocarpine-induced SE model of EPG, gliosis and neuronal death are progressing rapidly during the first week after SE (Fig. 6A,B). Data (***Brackhan et al., 2016***) suggests that the progression is slowing down during the second week after injury, reaching a plateau as indicated by a comparison of day 5 and day 14 post SE (Fig. 6). However, when looking at a longer time scale, a comparison of the immunoreactivity of neuron-specific nuclear protein (NeuN) in the hippocampus (***Zhang et al., 2015***) between days 7 and 60 suggests a profound progression of neuronal loss (Fig. 6C). Our model captures this temporal pattern of pathological development of gliosis and neuronal loss with a relative recovery after the acute neurological injury (Fig. 6A,B) and further progression of pathology in the chronic phase (Fig. 6C).

The modeling results suggest that despite the relative recovery of the BBB permeability, seizure burden grows due to the gradual increase in the degree of circuit remodeling (Fig. 6D). The pathological changes in circuity are happening in reaction to the remaining BBB leakage and ensuing damage to the neural population. The slowing of neuronal loss progression (Fig. 6B) and later progression during the chronic phase (Fig. 6C) are explained by the leveling off of the neuroinflammation after the initial injury and subsequent growth of neurotoxicity associated with the growth of seizure burden (Fig. 6D). Thus, seizures take over the propellent role in pathology development from the initial neurological injury. This results in the emergence of the complex temporal pattern of pathology progression in the pilocarpine animal model.

### Multicausality and degeneracy in EPG: Neuronal loss is sufficient but not necessary for inducing EPG

The role of neuronal loss in epilepsy development has been extensively debated. Massive neurodegeneration in the hippocampus, known as hippocampal sclerosis, is a common pathology of temporal lobe epilepsy and other epilepsy syndromes (***Thom, 2014***). Moreover, the extent of neuronal loss has been shown to be positively correlated with seizure frequency (***Lopim et al., 2016***). However, it is still an open question whether neuronal loss is a primary cause of EPG, its consequence, or both (***Tasch et al., 1999***; ***Kapur, 2003***; ***Sendrowski and Sobaniec, 2013***). In a study by ***Weissberg et al***. (***2015***), EPG with recurrent seizures was triggered in mice by induction of BBB disruption without evidence of neuronal loss. This indicates that neuronal loss may not be necessary for EPG. As discussed above, this phenomenon is readily explained by our model, which can produce EPG without cell death based on inflammation and BBB disruption alone (Fig. 3F). Given these results, we wondered if cell death, while not being *necessary* for EPG induction, may still be *sufficient* for it.

Indeed, our model predicts that neuronal loss alone can trigger the induction of EPG (Fig. 7A). Specifically, neuronal loss triggers slow remodeling of neural circuits, which gradually lowers the seizure threshold and increases the seizure rate. Mathematically, the presence of neuronal loss modifies the locations of the steady states (Fig. 7B): the ‘healthy’ steady-state and the unstable fixed point move towards each other, resulting in a non-zero seizure rate even when the system is resting in the ‘healthy’ steady state. Further increase of neuronal loss leads to a bifurcation where the ‘healthy’ steady state collides with the unstable fixed point at a certain value of neuronal cell loss *D*_critical_ ≈ 0.41 (Fig. 7B, for derivation see Appendix 7). For values of neuronal cell loss greater or equal than *D*_critical_, the development of progressive EPG towards the ‘epileptic’ steady-state is inevitable (Fig. 7A). In this case, the exact extent of neuronal loss determines the average time until the development of progressive EPG (Fig. 7C). However, progressive EPG is also possible in animals with a subcritical extent of neuronal loss due to stochasticity (*D* < *D*_critical_, see animals with D = 0.3 in Fig. 7A). The finding that neuronal loss is sufficient for EPG initiation and progression, while not being necessary for EPG in other types of injuries (Fig. 3), highlights the multicausal nature of EPG, where distinct processes may drive the process in isolation or in a convergent fashion. This is in line with a recent proposal that in degenerate systems (***Edelman and Gally, 2001***) multiple different pathological changes are sufficient but not necessary to cause hyperexcitability (***Ratté and Prescott, 2016***).

**Figure 7.**
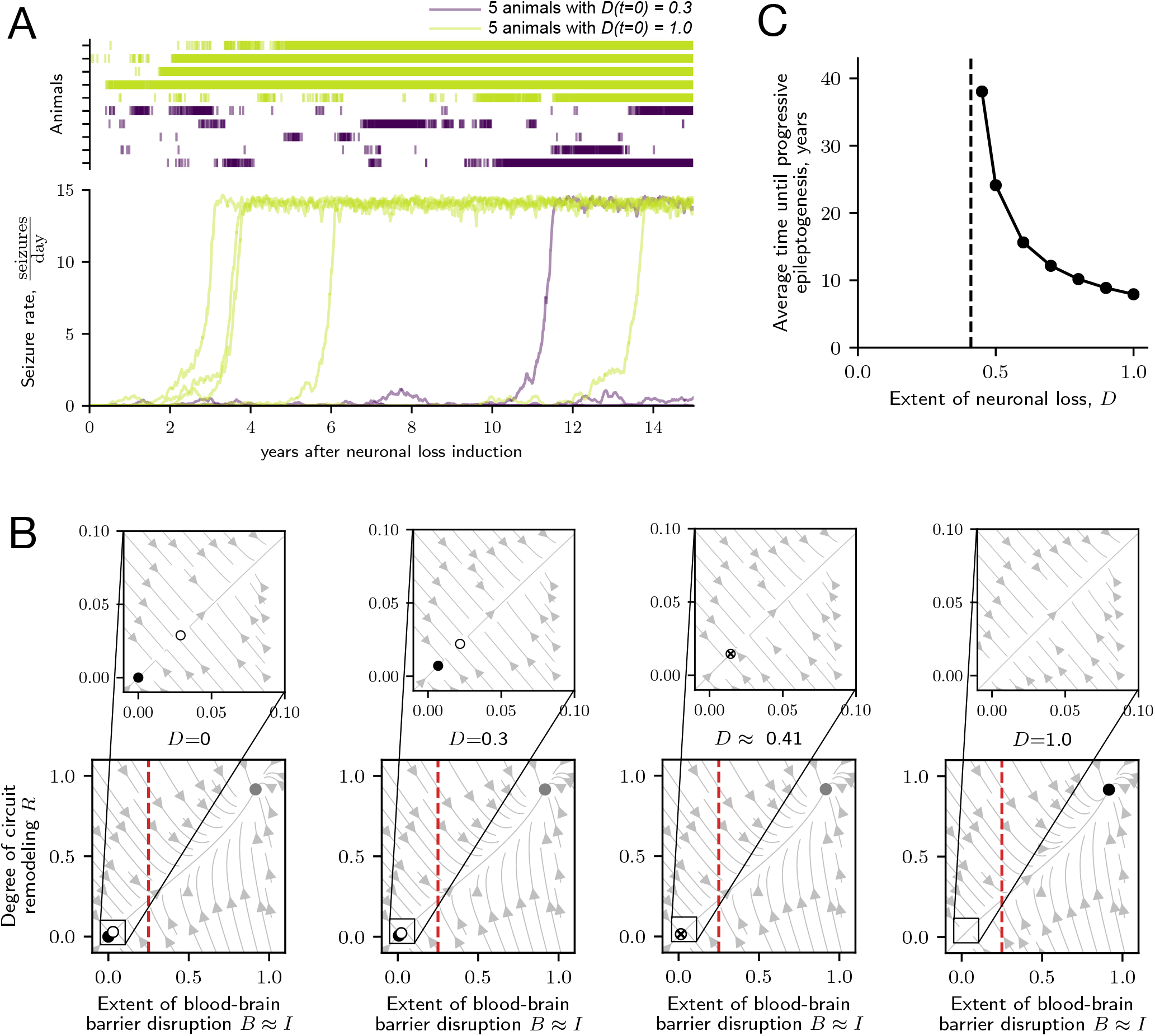
Model reveals that neuronal loss is sufficient but not necessary for EPG. **A**. EPG progression in 5 simulated animals with supercritical (*D* = 1.0 > *D*_critical_ ≈ 0.41) and subcritical (*D* = 0.3 < *D*_critical_ ≈ 0.41) extents of neuronal loss. The raster plots above seizure rate traces indicate seizure times of each animal. **B**. Effect of neuronal loss on the system stability illustrated with state space plots for rising extent of neuronal loss from left to right panels (*D* = 0; 0.3; ≈ 0.41; 1.0). Filled circles correspond to ‘healthy’ (black) and ‘epileptic’ (gray) steady states. Empty circles correspond to unstable fixed (saddle) points. Crossed empty circles correspond to semistable (one of the eigenvalues is zero) fixed points. For detailed analysis see Appendix 3. Red dashed line corresponds to neurotoxicity threshold Θ. **C**. Average time until inevitable progressive EPG for different extents of neuronal loss obtained with rate model. The time of progressive EPG was heuristically calculated as the time from the start of the simulation to the time point of the neuroinflammation *I*(*t*) reaching 90% of the value corresponding to the ‘epileptic’ steady state. Black dashed line corresponds to critical extent of neuronal loss *D*_critical_ ≈ 0.41.

### Simulation of therapeutic interventions reveals injury-specific targets and optimal time windows for treatment

Neuroimmune interactions are potential targets in the search for efficient treatments for pharmacoresistant epilepsy. However, not only selection of prominent targets for intervention, but also the timing and duration of interventions seems to matter. For example, application of rapamycin, which is suspected to have antiepileptogenic effect via restoration and strengthening of the BBB, over the period of 6 weeks after induced SE was efficient in the reduction of number of animals developing seizures, seizure frequency and extent of neuronal loss (***van Vliet et al., 2012***). In contrast, treatment that continued only for 2 weeks had no positive effect over a 6 week observation period (***Sliwa et al., 2012***). Our model provides an opportunity for simulating various intervention strategies, allowing for the selection of target and time window and exploring the effects of multi-target interventions.

Specifically, our model shows that a *permanent* suppression of the effect of seizures on BBB integrity (intervention I in Fig. 8A) prevents EPG in a simulated pilocarpine rodent model of epilepsy. The long term effect on seizure rate is illustrated in Fig. 8C. The extent of neuronal loss over the first half a week after injury does not differ among simulations with and without interventions (Fig. 8B). Figure 8D illustrates the impact of the suppression of the effect of seizures on BBB integrity. Under this suppression (right plot), only a single attractor corresponding to the ‘healthy’ steady state remains in the state space of the system.

**Figure 8.**
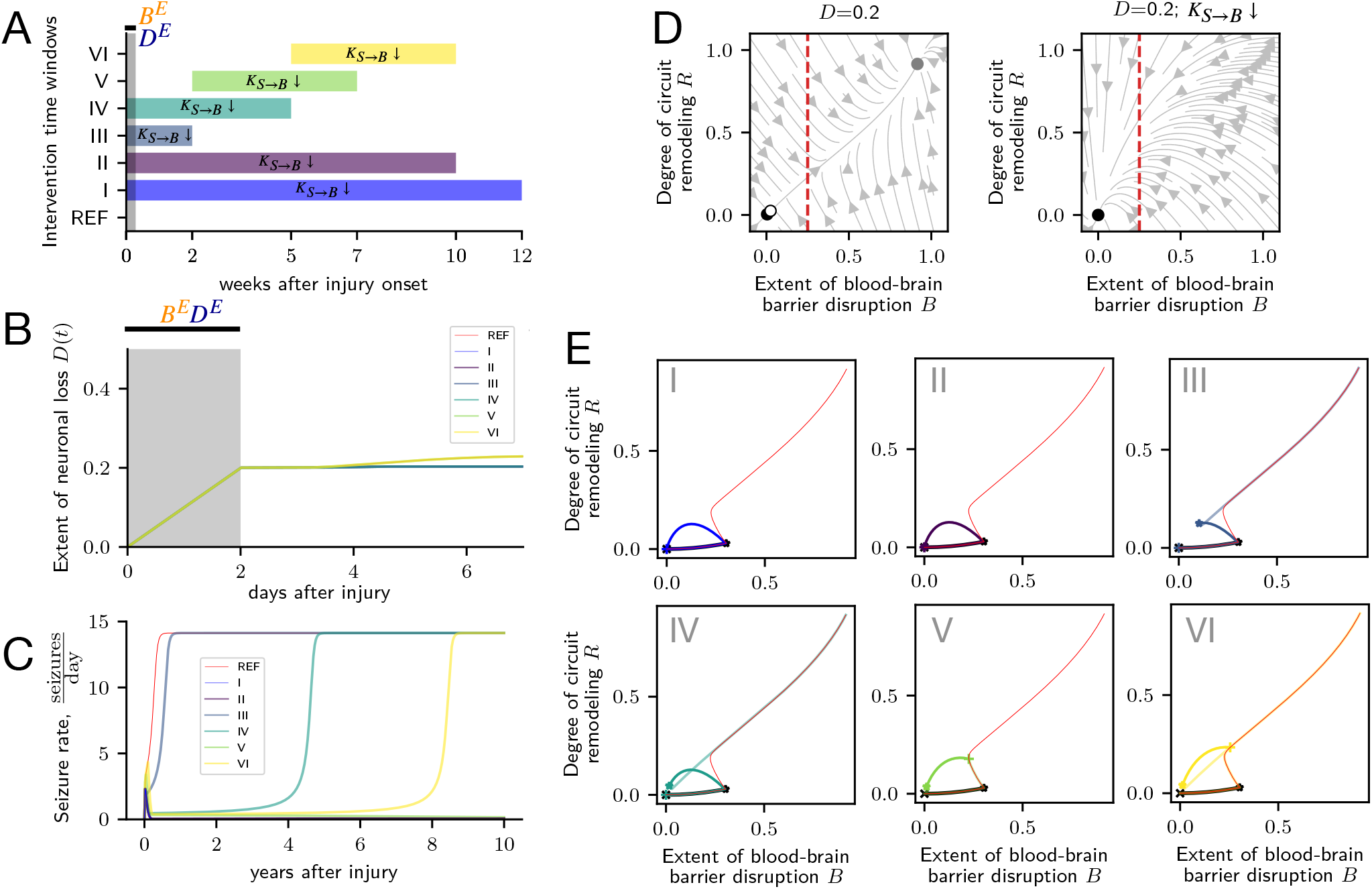
Modelling therapeutic intervention with suppression of seizure effect on BBB integrity reveals injury-specific time window for intervention in pilocarpine rodent model of epilepsy: **A**. Time windows for 6 various interventions and a reference simulation without intervention. Gray area corresponds to the time window of injury induction. The suppression of seizure effect on BBB integrity is simulated with 100-fold decrease of respective model variable *K*_*S*→*B*_ ↓. Simulations are performed using the rate model. **B**. Neuronal loss progression in animals exposed to various types of intervention. Gray area corresponds to the time window of injury induction. **C**. Seizure rate development in animals exposed to various types of intervention. **D**. State space plots illustrating the state of the system after the injury offset (D=0.2) without (left) and under the effect of intervention (*K*_*S*→*B*_ ↓, right). Filled circles correspond to ‘healthy’ and ‘epileptic’ steady states. Empty circle corresponds to the unstable fixed point. Red dashed line indicates the neurotoxicity threshold Θ. **E**. Response of the system to injury in animals exposed to various types of intervention illustrated in the B-R state space domain. Red lines correspond to the reference simulation without intervention. Solid black lines starting with ‘x’-symbol and ending with a star corresponds to the time interval of injury induction. Solid color lines starting with ‘x’-symbol and ending with a star correspond to time windows of intervention.

Further, we also investigated *transient* suppression of the effects of seizures on BBB integrity. Suppression during a long time window of 10 weeks (intervention II in Fig. 8A) is sufficient to prevent EPG (Fig. 8C,E), while a shorter window of 2 weeks (intervention III in Fig. 8A) does not suffice (Fig. 8C,E).

Interestingly, not only the duration of an intervention, but also its precise timing relative to the injury is crucial for successful prevention of EPG. For example, in the simulated pilocarpine rodent model of epilepsy, a suppression of the effects of seizures on BBB integrity for 5 weeks can prevent EPG when applied 2 weeks after the injury (intervention V in Fig. 8A). However, identical interventions starting at 0 or 5 week delays (interventions IV and VI in Fig. 8A) are inefficient (Fig. 8C,E).

Moreover, the time windows, during which interventions should be applied, have injury-specific durations and timings. For example, in the simulated TMEV infection rodent model, a 1 week long suppression of the effects of seizures on BBB integrity (intervention II in Fig. 9A) is sufficient to prevent EPG (Fig. 9C). Interestingly, in this model the intervention time window has to overlap with the time window of the injury effects. Interventions that are applied with 1 and 2 week delays (interventions III and IV in Fig. 9A) do not prevent EPG (Fig. 9C). In contrast to the simulation of the pilocarpine rodent model (Fig. 8B), in the TMEV infection model interventions that are applied during and after the period of injury effects lead to different levels of neuronal loss after the injury offset (Fig. 9B). This results in different structures of the systems’ state spaces: in case of interventions I and II, which overlap with the injury time window (Fig. 9A), a lower extent on neuronal loss leads to preservation of a larger basin of attraction of the ‘healthy’ steady state (Fig. 9D). This is in contrast to interventions III and IV (Fig. 9A), for which the ‘healthy’ steady state and the separatrix are in close proximity (Fig. 9E). Therefore, interventions that are not applied during the first week after injury onset are insufficient for the prevention of EPG (Fig. 9C).

**Figure 9.**
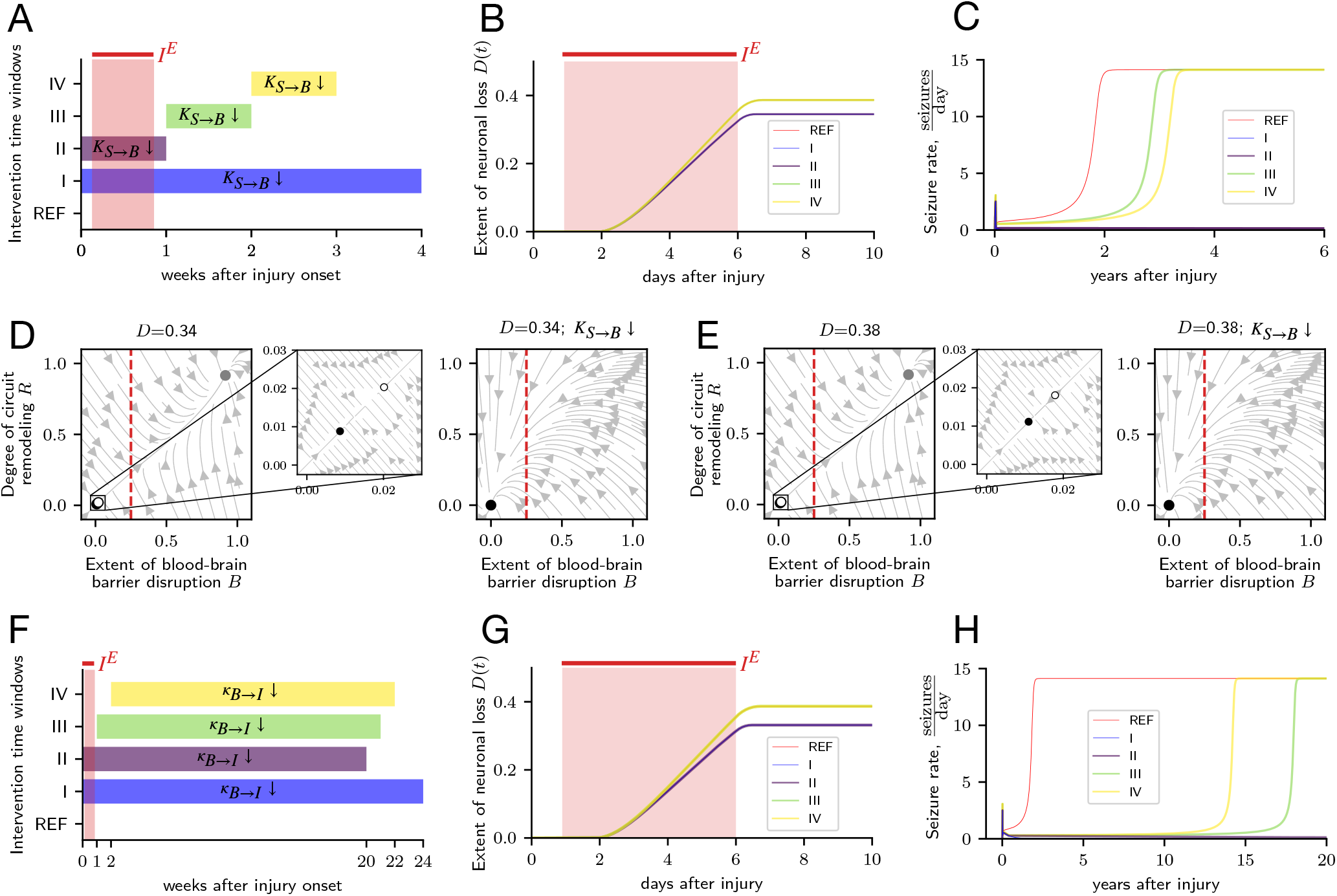
Modelling therapeutic interventions suppressing the effects of seizures on BBB integrity and activation of glia by factors infiltrating the parenchyma reveals injury-specific optimal time windows for intervention in TMEV infection rodent model of epilepsy: **A**. Time windows of 4 interventions suppressing the effects of seizures on BBB integrity. REF indicates reference without intervention. Light red area corresponds to the period of injury induction. The suppression of the effect of seizures on BBB integrity is simulated with a 100-fold decrease of the respective model variable *K*_*S*→*B*_. Simulations are performed using the rate model. **B**. Neuronal loss progression for the different time windows of *K*_*S*→*B*_ reduction. Light red area corresponds to the period of injury induction. **C**. Seizure rate development for different intervention time windows. **D**. State space plots illustrating the state of the system after the injury offset (*D* = 0.34) for interventions that coincided with injury time window without (left) and under the effect of intervention (right). Filled circles correspond to ‘healthy’ and ‘epileptic’ steady states. Empty circle corresponds to the unstable fixed point. Red dashed line corresponds to neurotoxicity threshold Θ. **E**. Same as **D** but for a higher value of *D* = 0.38. **F**. Time windows of 4 interventions with suppression of activation of glia by factors infiltrating the parenchyma. Light red area corresponds to the period of injury induction. The suppression of activation of glia by factors infiltrating the parenchyma is simulated with 100-fold decrease of the respective model variable *κ*_*B*→*I*_. Simulations are performed using the rate model. **G**. Neuronal loss progression for different time windows of *κ*_*B*→*I*_ reduction. Light red area corresponds to the time window of injury induction. **H**. Seizure rate development for different time windows of *κ*_*B*→*I*_ reduction.

In a next step, we were interested in investigating the effects of different types of interventions in the TMEV infection model of epileptogenesis. Interestingly, the necessity of early intervention already during the active injury effect on the system also applies to the TMEV infection model. Here, we have simulated an intervention that suppresses glial ability to be activated by infiltrating blood factors with a 100-fold decrease of the respective model variable *κ*_*B*→*I*_ ↓ (Fig. 9F). This type of intervention requires a much longer time window of application, but also requires application during the first week after injury onset (Fig. 9F-H). In sum, our model can be used as a framework for simulation of intervention strategies. It provides means for studying the efficiency of various therapeutic targets, intensities and time intervals of intervention in an injury-specific manner.

## Discussion

The pathophysiology of EPG is associated with the activation of innate and adaptive immune responses, disruption of BBB integrity, neuronal loss, circuit remodeling and various other processes acting over a range of different timescales. Furthermore, the injury-specific time courses of clinical markers point to a bewildering complexity of the disease. This makes understanding EPG and the development of effective treatments a formidable challenge. Computational and mathematical modeling can be a valuable tool in understanding such complex systems. Indeed, other epilepsy-associated phenomena, such as ictogenesis, have already been successfully studied with mathematical modeling methods (***Jirsa et al., 2014***; ***Proix et al., 2017***; ***Jirsa et al., 2017***). Here, we have presented the first-of-its-kind mathematical model of EPG in the context of acquired epilepsy. Our model explains a wide range of EPG phenomena and is a tool for testing different interventions *in silico*, while generating testable predictions regarding their effectiveness. The model describes the interaction between neuroinflammation, BBB disruption, neuronal loss, circuit remodeling, and seizures in response to neurological injury. Mathematically, the model consists of a system of coupled stochastic non-linear ordinary differential equations. Our formal analysis of the model has revealed the existence of two stable fixed points, representing the healthy state and the state of a developed epilepsy.

Our model explains how EPG is triggered by very different types of neurological injuries. Here, we have focused on three such injuries: infection as represented by a TMEV rodent model; chemical intoxication as represented by pilocarpine SE rodent model; and BBB leakage as represented by BBB disruption rodent model. We have found our model to be in good agreement with the experimental data from these animal models using a single set of parameters for all simulations.

The model captures injury-specific characteristics of EPG such as temporal patterns of seizure occurrence, the progression of neuronal loss, neuroinflammation and BBB disruption. Interestingly, the model explains long timescales (years and decades) of disease development despite time-limited injuries that directly affect the central nervous system for much shorter durations (days). Mathematically, these unexpectedly slow timescales of EPG are explained by a slowing of the system’s dynamics in the vicinity of an unstable fixed point. This resembles the emergence of slow dynamics in, e.g., wound healing after injury with paradoxically long scar formation (***Adler et al., 2020***). Moreover, our model describes the dependence of the latent period duration, the seizure burden, and the risk of EPG on the intensity of an injury — the dose-dependence effects of injury intensity observed in various human and animal models of EPG.

Our model also captures the multicausal nature of epilepsy. For example, our model explains how, on the one hand, neuronal loss alone may be *sufficient* to induce EPG, but, on the other hand, it is not at all *necessary* for EPG. This is in agreement with a recent observation that, in neuronal systems with degenerate mechanisms, several distinct pathologies are sufficient but not necessary to account for the hyperexcitability (***Ratté and Prescott, 2016***). Furthermore, our model suggests that the variability of EPG outcomes originates in part from the stochastic nature of epileptic seizures, which can push the system from the basin of attraction of the ‘healthy’ steady state to that of a developed epilepsy.

In order to demonstrate the utility of the model for generating testable predictions of therapeutic interventions, we performed simulations with different intervention targets and time windows for different initial injuries. Our results suggest that therapeutic interventions applied during only a short but critical time window may be just as effective as long-term interventions. Moreover, the optimal time windows for interventions are injury-specific. For example, in the case of a TMEV infection model, the intervention has to be applied during the first week after injury onset and without a delay. In the pilocarpine model, in contrast, a 5-week intervention starting after 2 weeks prevented EPG, while earlier or later interventions were ineffective.

Due to its simplicity, our model also has a number of important limitations. For example, the complex process of neuroinflammation is described by just a single “coarse-grained” variable. This aids mathematical analysis, but it complicates the interpretation of simulation results. Furthermore, the model does not distinguish different seizures types (e.g. focal vs generalized) nor does it allow for an evolution of seizure severity and duration throughout EPG. Besides processes described by the model, phenomena such as channelopathies, neurogenesis, gene transcription, epigenetic modifications, and others are also associated with EPG, but yet to be accounted for in our modeling framework.

Also, the model does not include any positive antiepileptic aspects of neuroinflammation, i.e. processes aimed at the maintenance of healthy central nervous system function. Future extensions of the model should take into account such protective aspects of neuroimmune interactions. This will allow for computational modeling of paradoxical phenomena such as inflammatory preconditioning and epileptic tolerance. These, together with further exploration of injury-specific targets and time windows for therapeutic interventions, are promising directions for future work. Finally, while we view it as a strength of the model that it explains data from different animal models with a single set of parameters, we acknowledge that there is always inter-individual variability of physiologic parameters. It will therefore be interesting to investigate how parameter variations change the individual susceptibility to EPG and its trajectory. Such an understanding will facilitate the development of individualized interventions in the spirit of precision medicine.

## Acknowledgments

The work was supported by the grant LOEWE CePTER – Center for Personalized Translational Epilepsy Research (to DB, FL, PJ and JT). DB is supported by the International Max Planck Research School (IMPRS) for Neural Circuits. FL is supported by the Swartz fellowship in Theoretical Neuroscience at University of Washington in Seattle. AV is supported by the Era-Net Neuron Ebio2 consortium. PJ is supported by the BMBF grant (No. 031L0229) and the funds from the von Behring Röntgen Foundation. JT is supported by the Johanna Quandt Foundation.

## Appendix 1

This figure illustrates the sigmoid function of seizure rate dependence on seizure promoting effects (Eq.2). This function is used in simulation and vizualization of simulation outcomes.

**Appendix 1 Figure 1.**
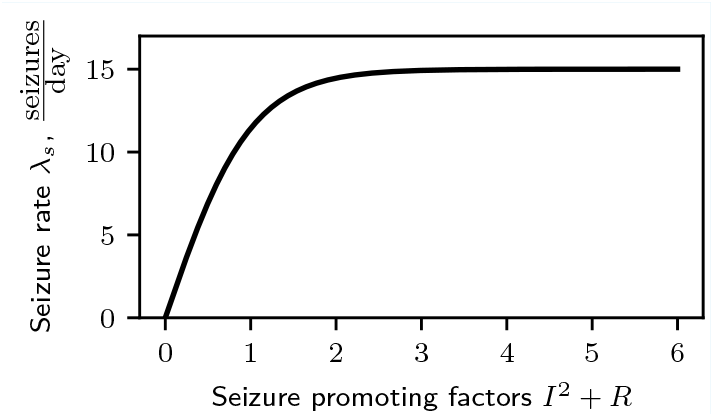
Seizure rate dependence on seizure promoting factors: neuroinflammation and circuit remodeling.

## Appendix 2

## Timescale separation

In this work, we assume that the process of neuroinflammatory reaction evolves faster than BBB disruption and recovery of its integrity, neuronal loss, and circuit remodeling: *τ*_*I*_ ≪ *τ*_*B*_, *τ*_*D*_, *τ*_*R*_. Thus, under condition of absence of the neuroinflammatory external input *I*_*E*_, we can perform a time scale separation, which at equilibrium will result in *I* ≈ *κ*_*B*→*I*_*B*, and the system described in Eq. 1 becomes:

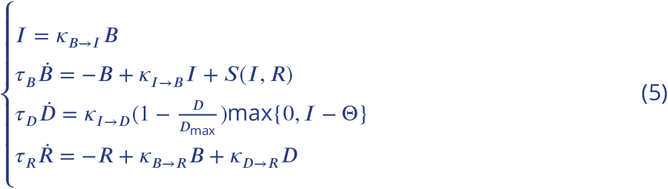

From Eq. 5, we can obtain a system of equations for fixed values of neuronal loss extent *D* = *D*_const_, where 0 ≤ *D*_const_ ≤ *D*_max_. The resulting system of equations describes the system in the dynamical regimes characterized by the absence of neurotoxicity *I* ≈ *κ*_*B*→*I*_ *B* < Θ:

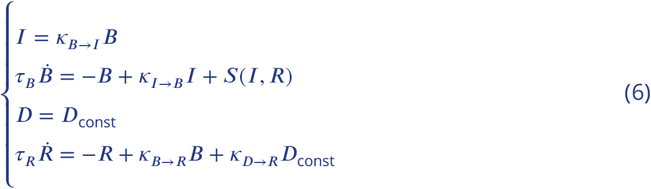

Substituting *I* = *κ*_*B*→*I*_ *B* in the equation for the extent of BBB disruption, we obtain the system described in *B* − *R* dimensions. It is used for analysis and visualization of dynamics with state space plots for variables *B* and *R*:

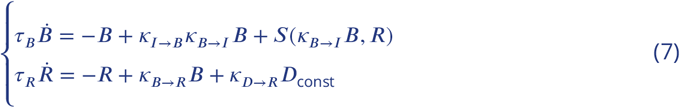

where *I* = *κ*_*B*→*I*_ *B* < Θ and *D* = *D*_const_.

## Appendix 3

## Stability analysis

In this section, we are going to analyse the stability of the steady states of the system (state space composition).

In a steady state, 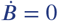 and 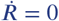. From Eq. 7 we obtain:

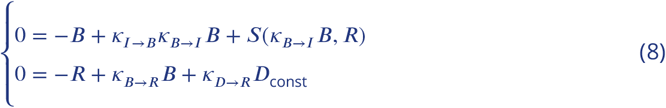

Substituting *S*(*I, R*) from Eg. 4:

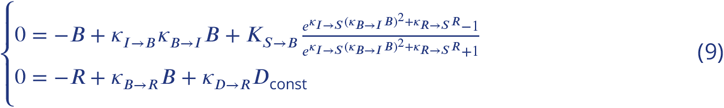

The steady states (fixed points) in the system are the result of intersection of 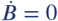 and 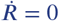, which gives us (inserting the parameter values from Appendix 4) two equations:

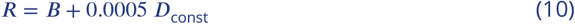

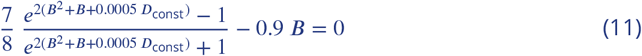

The intersection of the latter Eq. 11 with the horizontal axis will give alll *B*\* that satisfy 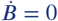 and 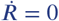. And corresponding values of *R*\* can be found using Eq. 10.

The Jacobian of the linearized system around each fixed point is:

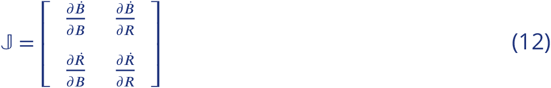

where 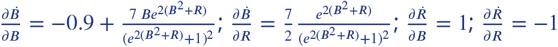.

The eigenvalues for each fixed point [*B*\*, *R*\*] are the eigenvalues of the Jacobian evaluated at [*B*\*, *R*\*]. Analysing the fixed point positions and their corresponding eigenvalues, we can describe the system state space.

1. when 0 ≤ *D*_const_ < *D*_critical_ the system has 3 fixed points (Fig. A3.1): a stable steady state (negative eigenvalues) around the origin; a saddle point (one positive and one negative eigenvalue); a stable steady state (negative eigenvalues) distanced from the origin in the first quadrant;
2. when *D*_const_ = *D*_critical_ ≈ 0.41 the system undergoes the saddle node bifurcation and has 2 fixed points (Fig. A3.1): a stable steady state (negative eigenvalues) distanced from the origin in the first quadrant, and a semistable point (one eigenvalue equal to 0) in the position of collision of two fixed points;
3. when *D*_critical_ < *D*_const_ ≤ *D*_max_ the system has 1 fixed point - a stable steady state (negative eigenvalues) distanced from the origin in the first quadrant (Fig. A3.1).

**Appendix 3 Figure 1.**
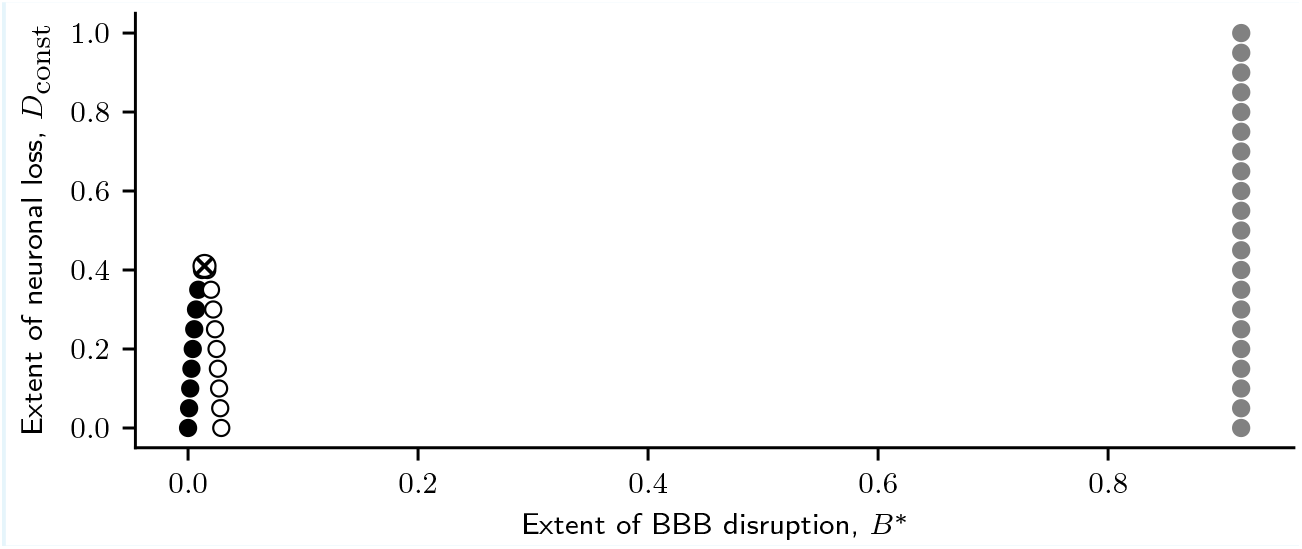
Saddle node bifurcation illustrated with collision (crossed circle) of stable (black circles) and unstable (white circles) steady states at critical value of extent of neuronal loss (*D*_const_ = *D*_critical_ ≈ 0.41). The third fixed point (gray circles) shows low sensitivity to change of neuronal loss due to low value of *κ*_*D*→*R*_. For values *D*_const_ > *D*_critical_, only one stable steady state (gray circles) exist.

The code for numerical calculation of the fixed point values of *B*\* and *R*\*, and the eigenvalues used in the stability analysis can be found at https://github.com/danylodanylo/math-model-epileptogenesis.git. For calculation of bifurcation parameter value *D*_critical_ (the extent of neuronal loss at which the saddle node bifurcation is occurring) see Appendix 7.

## Appendix 4

**Appendix 4 Table 1.**
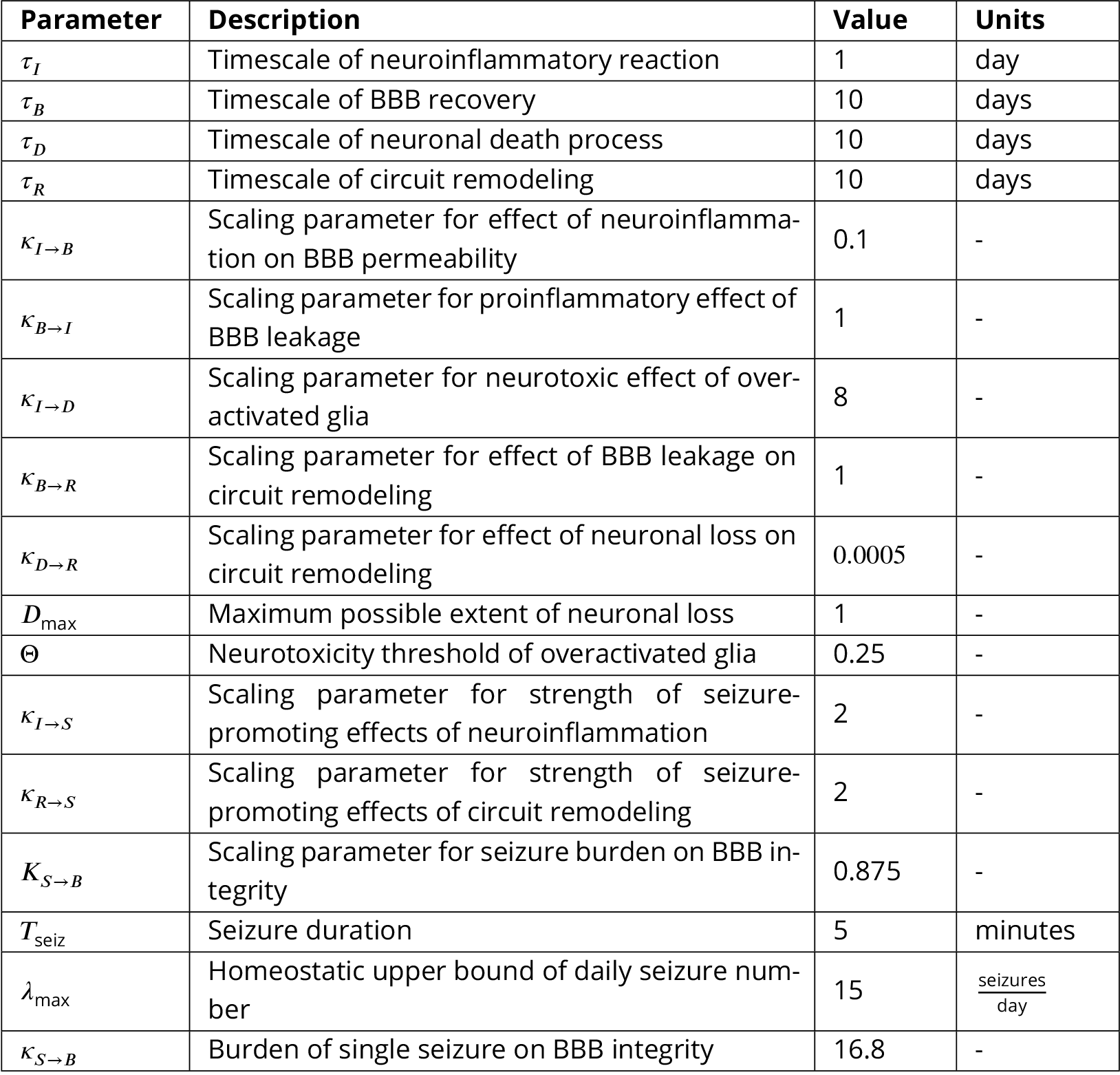
Model parameter descriptions and values.

## Appendix 5

**Appendix 5 Table 1.**
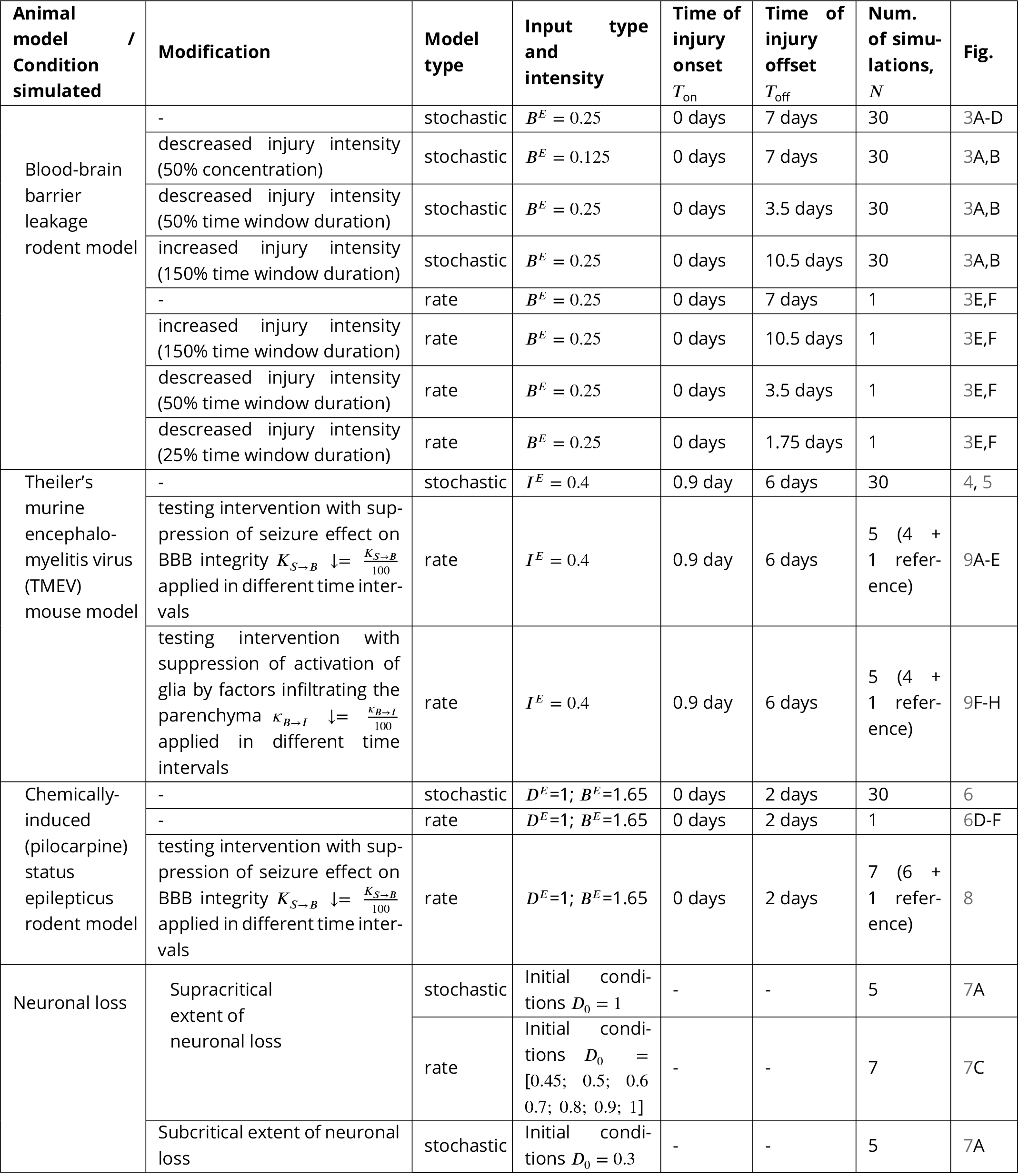
Detailed specifications of performed simulations.

## Appendix 6

## BBB disruption rodent model data used in Fig. 3A-B

Following data from ***Weissberg et al***. (***2015***) were used in this study:

- latent period duration of 4.9 ± 1.3 days (mean ± SEM), N=10;
- spontaneous seizures frequency of 1.16 ± 0.16 seizures per day (mean ± SEM), N=10.

## Theiler’s murine encephalomyelitis virus (TMEV) mouse model data used in Fig. 4A-C

Following data from ***Patel et al***. (***2017***) were used in this study:

- number of seizures per day for N=11 mice was extracted from Figure 2 (***Patel et al., 2017***). Average seizure frequency per mice was calculated for 3 time intervals: day 1 post infection, days 2-7 post infection and days 8-15 post infection;
- TNF protein fold change (relative to PBS-injected control mice) on day 1 post infection (N=8): 6.9 ± 0.6, day 5 post infection (N=6): 206.2 ± 14.9, day 14 post infection (N=5): 34.8 ± 7.1. Data presented in mean ± SEM.

Following data from ***Kirkman et al***. (***2010***) were used in this study:

- neuronal cell loss score for 2 hippocampi (mean ± SEM) on days 1-35 post infection from Figure 2 (***Kirkman et al., 2010***), N=4-13 per time point group.

## Chemically-induced (pilocarpine) SE rodent model data used in Fig. 6A-C

Following data from ***Brackhan et al***. (***2016***) were used in this study:

- microglial activation score for the hippocampus (mean ± SEM) on days 0, 2, 5, 14 post SE from Figure 4 (***Brackhan et al., 2016***), N=3-5 per time point group;
- neuronal cell loss score for the hippocampus (mean ± SEM) on days 0, 2, 5, 14 post SE from Figure 4 (***Brackhan et al., 2016***), N=3-5 per time point group.

Following data from ***Zhang et al***. (***2015***) were used in this study:

- NeuN-immunoreactive cells count per mm^2^ in the hippocampus of pilocarpine treated animals from Figure 5 (***Zhang et al., 2015***). Fraction of cells missing (in %) was computed for days 7 and 60 after pilocarpine injection relatively to values for untreated animals.

## Appendix 7

## Estimation of critical extent of neuronal loss

In this section, we are going to calculate the critical extent of neuronal loss *D* = *D*_const_ = *D*_critical_, at which ‘healthy’ steady state collides with the unstable fixed point and only one stable steady state remains for *D* > *D*_critical_. From equations describing system state for fixed extent of neuronal loss (Eq. 6) and seizure burden function (Eq. 4), we derive steady state equation for BBB disruption:

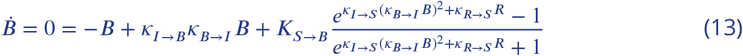

and steady state equation for circuit remodeling:

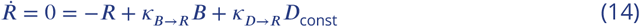

from which we derive:

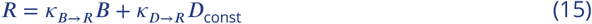

which will be referred to as *linear R*.

Inserting the parameter values (Appendix 4) into Eq. 13:

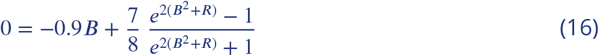

Defining 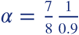, we can derive *B* from Eq. 16:

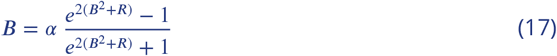

Defining 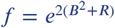 from Eq. 17, we obtain:

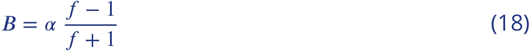

From Eq. 18, we can derive:

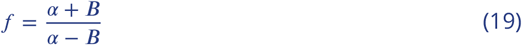

Now, we replace *f* with 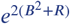:

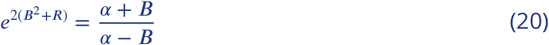

Taking logarithms of both sides:

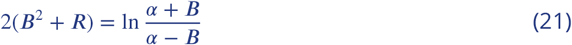

From Eq. 21, we can obtain *nonlinear R* equation:

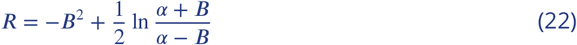

The intersections between *linear R* and *nonlinear R* will give us all the fixed points of the system. With the parameters defined in Appendix 4, this system of equations always has at least one fixed point for *B* < 1. In addition to this fixed point, a saddle node bifurcation can emerge when two additional fixed points are generated as a result of a change of parameter in the equations. Assuming that *D* could play the role of such a bifurcation parameter, we need to find its critical value such that *linear R* becomes tangential to *nonlinear R*. Decreasing this critical value would result in the emergence of two fixed points; however, increasing this value beyond *D*_critical_ would result in no intersection between the nullclines, and hence the system will have only one fixed point which was defined before. To find the value of *D* = *D*_critical_ in *linear R* which results in a tangent line to the nonlinear curve, firstly, we need to find the first derivative of *nonlinear R* with respect to *B*. Secondly, we should find all those values *B*\* = *s*, where *s* is equal to the slope of *linear R*, or in other words, 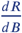 in *linear R* that is equal to *κ*_*B*→*R*_ = 1 (Eq. 15, Appendix 4). This indicates that we should find *B*\* in 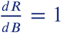 for the *nonlinear R*. Finding the first derivative of Eq. 22:

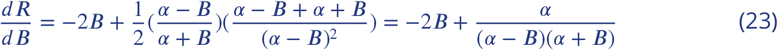

Equating Eq. 23 to *κ*_*B*→*R*_ = 1, we obtain a polynomial:

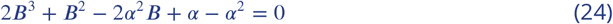

Solving this polynomial numerically (code available at https://github.com/danylodanylo/math-model-epileptogenesis.git), we get the following values for *B*\*= [−1.259; 0.7448; 0.01439]. Since BBB disruption variable can not take negative values, the solution [−1.259] is discarded. Using equation for *nonlinear R* (Eq. 22), we can calculate the corresponding *R* values for *B*\*= [0.7448; 0.0143]:

*R*\*=[0.4560; 0.0146].

Note that these values of *R*\* should hold in both *nonlinear R* and *linear R*, since they are the result of intersections between the nullclines. From *linear R* (Eq. 15), we can derive the equation for *D*_const_:

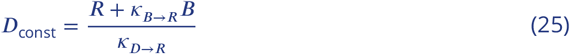

For values of *B*\* and *R*\*, we can calculate *D*\*_const_=[−577.5155; 0.4103]. Neuronal loss extent can not take negative values, so we have to discard one of the solutions [−577.5155]. Thus, we have found the only critical extent value of neuronal loss *D*_critical_ = 0.4103 ≈ 0.41.

---

**Figure 4-Figure supplement 1.**
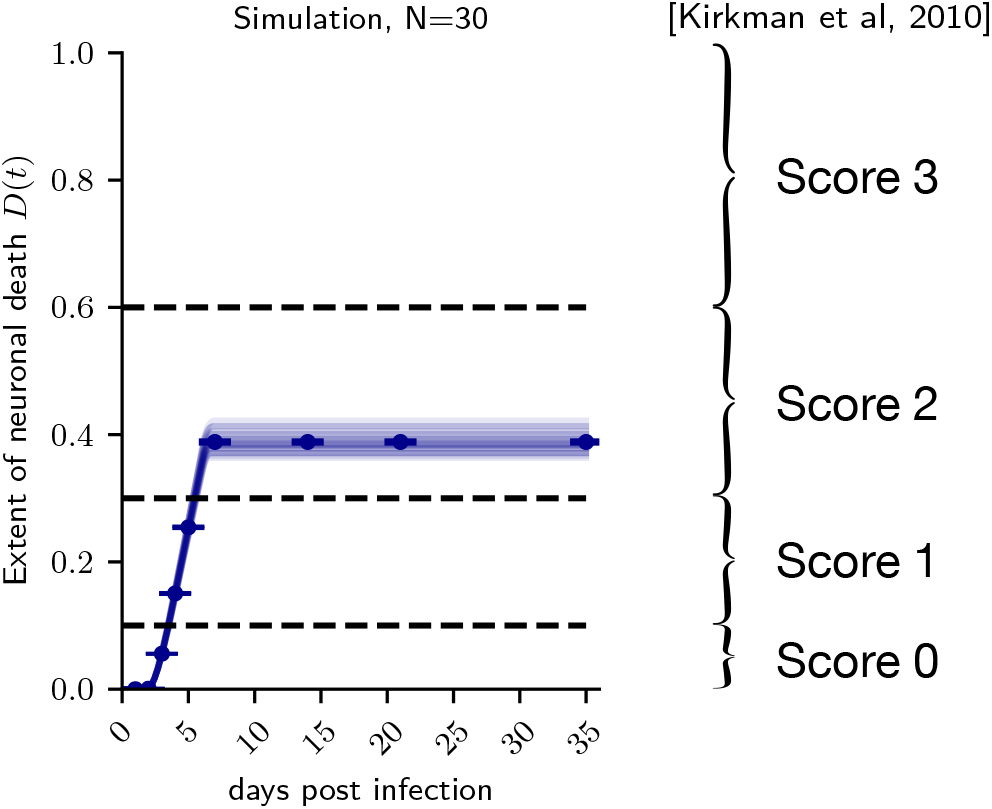
Neuronal loss score computation (masking procedure) from *Kirkman et al*. (*2010*): Raw neuronal death data from TMEV model simulation (left) and neuronal loss score computation scheme (right). Horizontal dashed lines on the left correspond to 10%, 30% and 60% extent of neuronal loss, which are the border values separating score values in the scheme from *Kirkman et al*. (*2010*). In *Kirkman et al*. (*2010*), neuronal loss score data are presented as a sum of scores for 2 hippocampi (maximum score: 3 × 2 = 6). Thus, neuronal loss score computed for simulated TMEV animals was multiplied by factor of 2 for comparability with experimental data. Absence of variability (0 SEM) in Fig. 4C is explained by ‘masking out’ of variability in neuronal loss score computation (left).

